# Sex-specific multigenerational epigenetic responses to real-world chemical mixture exposure in an outbred sheep model

**DOI:** 10.64898/2026.04.08.717152

**Authors:** Oliver G Hargreaves, Wing Yee Kwong, Andrew Warry, Desmond AR Tutt, Vasantha Padmanabhan, Neil P Evans, Richard G Lea, Michelle Bellingham, Kevin D Sinclair

## Abstract

Establishing whether real-world environmental chemical (EC) exposure can induce heritable epigenetic modifications in large, outbred mammals is key to determining long-term developmental impacts of the human exposome. Using an established biosolids-treated pasture (BS) sheep model, we investigated whether gestational exposure to low-level mixtures of EC induced heritable changes in DNA methylation across three generations of sheep. Reduced-representation bisulfite sequencing of liver, blood, and sperm, combined with a structured, lineage-controlled breeding design, revealed widespread but lineage- and sex-specific differentially methylated loci (DML) in F1 offspring, with detectable alterations evident in F2 and F3 descendants. Although most DML were unique to individual sire lineages, or to a single generation, subsets of loci showed repeated involvement across generations and were associated with altered gene expression in F3 descendants. Sperm from F1 males exhibited reduced methylation at numerous loci and, together with seminal plasma, revealed differential expression of several microRNAs. These effects, however, showed limited persistence in F2 males, indicative of intergenerational rather than fully transgenerational persistence. Collectively, these findings demonstrate that complex, low-level chemical exposures can elicit recurrent, sexually dimorphic epigenetic responses in outbred species, but underscore the challenge of disentangling exposure-induced inheritance from genetically regulated methylation variation.

**Significance Statement:** Environmental chemical (EC) exposures are ubiquitous, yet their capacity to induce heritable epigenetic changes in large, genetically diverse mammals is poorly understood. Using a real-world exposome-based sheep model, we demonstrate that low-level gestational EC exposure leads to sexually-dimorphic and lineage-dependent alterations in DNA methylation that can extend to unexposed descendants. Although genetic ancestry exerts a dominant influence over these responses, repeated alterations at specific loci suggests that environmentally induced epimutations can reoccur across generations in certain genomic contexts.

## Introduction

Exposure to complex mixtures of environmental chemicals (EC) prior to and throughout pregnancy is of concern in developmental toxicology, particularly due to the potential for long-lasting effects on offspring reproductive and metabolic health (1,2). In this regard, epigenetic modifications to DNA methylation provide a compelling mechanism by which such exposures can exert their influence, with the potential to extend across generations (3,4). The extent to which low-level EC exposure during gestation can alter the epigenome in large, precocial outbred mammals like humans, and whether these modifications can be inherited, remains unknown. A powerful model for studying these issues involves grazing pregnant sheep on pastures treated with biosolids (BS), a processed by-product of human sewage treatment that is used globally as an agricultural fertilizer (5). Biosolids contain low concentrations of a mixture of anthropogenic contaminants (including phthalates, per- and polyfluoroalkyl substances, flame retardants, pharmaceuticals and organochloride pesticides), rendering this model representative of the human exposome both in terms of composition and concentration of EC (6).

Early studies using the BS-sheep model revealed a range of adverse effects on the fetal and adult hypothalamic-pituitary-gonadal axis (7-11). More recent BS studies working with the cohort of sheep reported in the current article describe a testicular dysgenesis-like phenotype in F1 juvenile and adult male offspring similar to that reported in humans (12-14), effects on puberty onset in both sexes (15) and coincident evidence of reduced ovarian reserve in juvenile females (16,17). Metabolic studies have also demonstrated dysregulation in both maternal (F0) and adult F1-offspring, with evidence of inter-individual variability in responses to EC exposure (18-21).

To date, there has been no investigation of the epigenetic mechanisms that could underly these phenotypic effects within the BS model, or in large animal models of EC exposure more generally (1,22). As such, it is unknown if gestational exposure to EC from BS induces DNA methylation changes in offspring and whether such effects can be transmitted to later non-exposed generations. Elucidation of such heritable epigenetic changes in this model would be informative, given the physiological similarities between sheep and humans, and the outbred, genetically diverse nature of sheep as a model species. Evidence of heritable changes in DNA methylation following exposure to EC is limited currently to studies with rodents and to single or small mixtures of chemicals (e.g., vinclozolin, Bisphenol A, phthalates) (3,23). The investigation of inheritance of EC-induced epigenetic marks in an outbred mammalian species, however, presents challenges. Firstly, genetic variation within a population can confound the attribution of inherited differences in DNA methylation to exposure rather than underlying genotype (24,25). Furthermore, DNA methylation undergoes two rounds of global reprogramming during early embryogenesis and gametogenesis (26), rendering stable transmission of acquired marks inherently difficult (27). Finally, sexual dimorphism in epigenetic responses following EC exposure (28,29) must also be factored into any analysis.

With the foregoing discussion in mind, the present study sought to determine if gestational exposure to a real-life mixture of EC using the BS-sheep model would induce heritable changes in DNA methylation that would span three generations of offspring and include the third (F3) non-exposed generation. Reduced representation bisulfite sequencing (RRBS) was performed on descendant DNA extracted from liver (F1 to F3), sperm (F1 and F2) and blood (F2). In addition, RNA sequencing, with a focus on microRNA, was performed on sperm and seminal plasma from F1 and F2 male offspring, to provide an insight into putative ‘reconstructive’ modes of male germline epigenetic inheritance (27,30). Consideration was given throughout to the breeding structure of each of the three descendent populations to (i) standardize genetic variation between treatment groups within generation, (ii) avoid family matings and (iii) facilitate the tracking of differentially methylated loci (DML) across generations.

## Results

### Fertility and pregnancy outcomes

The breeding schedule devised for this study (Fig. S1) generated genetically matched (across the two experimental treatments) family groups for generations F1, F2 and F3. There was no effect of BS exposure of F0 ewes during the 6 weeks leading up to AI and gestation on pregnancy establishment, litter size, term delivery and sex distribution of offspring (Tables S1, S4). The same was true for F1 and F2 inseminations leading to the birth of F2 and F3 offspring (Tables S2, S3, S5, S6).

### F1 epigenetic effects

#### Liver DNA methylation

Samples were collected 8 weeks after birth from 16 F1 male and female offspring representing the four half-sib family groups for each of the two treatments (Table S4). RRBS reads covered 13,605,795 methylated cytosines, consisting of 50.9% CpG, 1.9% CHH and 2.3% CHG (a distribution repeated across all subsequent generations). Hierarchical clustering of methylated cytosines revealed a strong paternal (sire) influence within which were nested the effects of offspring sex and treatment respectively (Fig. 1A). Subsequent analyses of differentially methylated loci (i.e., CpGs ≥15% difference (DML)) were performed in two sets, one split by sire (differences averaged across sex) and one split by sex (differences averaged across sire). When split by sire, 25,374 unique DML were observed, 93.4% of which were specific to a single sire (Fig. 1B, F). When split by sex, 3,725 DML were observed with 98.9% specific to a single sex (Fig. 1C, F). Mean methylation differences between treatments, averaged across sire, were similar quantitatively for male (6.9% net increase in BS exposed; Fig. 1D) and female (7.4% net increase in BS; Fig 1E) offspring. Further analyses identified 502 unique genes containing genic (i.e., from 3 kb upstream of the TSS to 3 kb downstream of the 3’UTR) DML, 42.1% of which were specific to males and 54.5% specific to females (File S1A, B).

**Figure 1.**
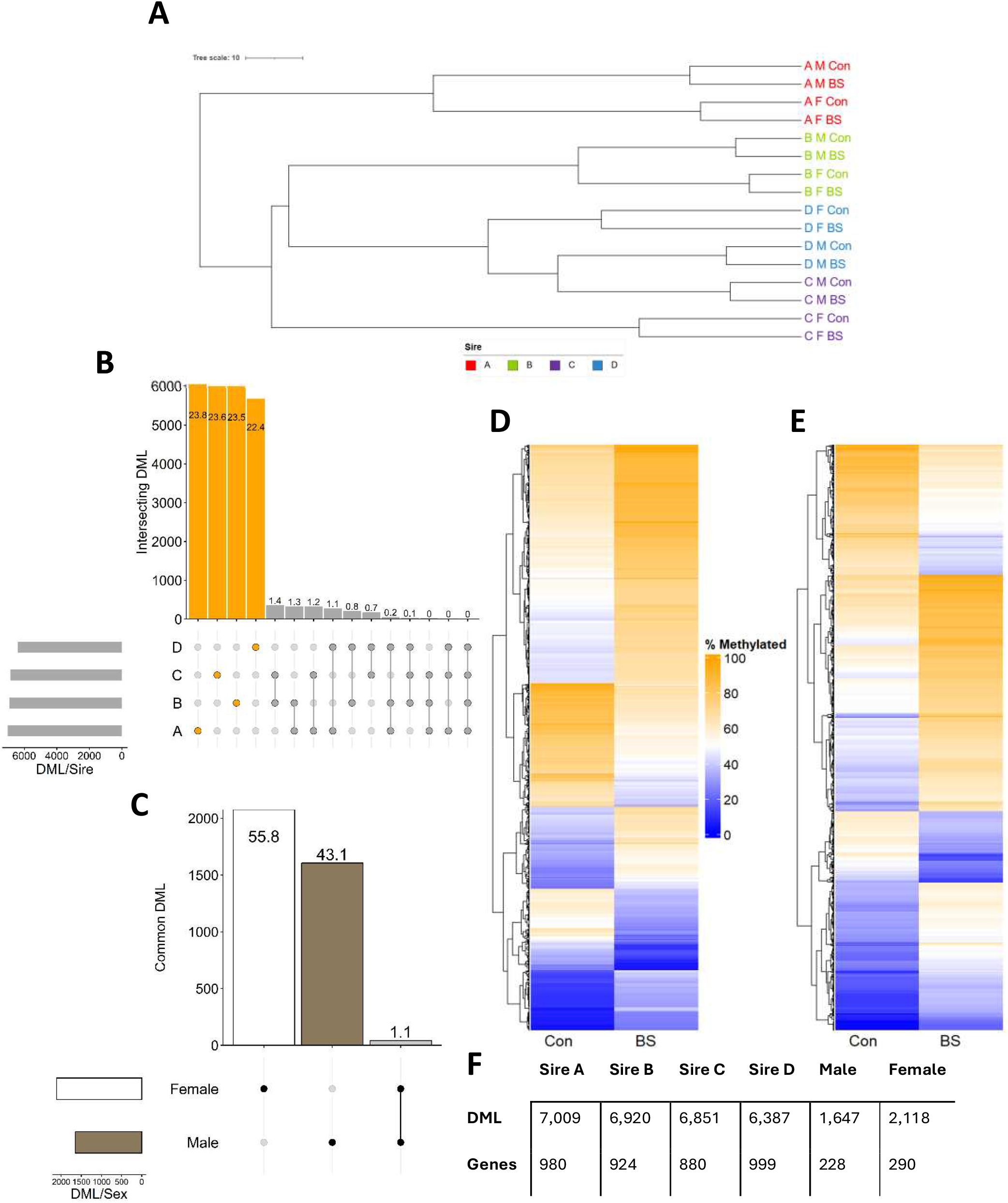
Methylation characteristics of F1 liver. **A.** Hierarchical clustering of DNA methylation in F1 liver samples. Distances calculated from top 3 principal components. Label colors indicate sire groupings. **B**. Upset plot of overlapping differentially methylated loci (DML) (15% threshold) across the four F1 sire groups (Orange: sire specific, Grey: overlapping; values on bars denote percentage of total DML). **C**. Upset plot of overlapping DML across male and female F1 livers (Black/White: sex specific, Grey: overlapping, values on bars denote percentage of total DML). **D**. Heatmap highlighting mean methylation of DML in males in the Control (Con) and Biosolids (BS) exposed experimental groups. **E**. Heatmap highlighting mean methylation of DML in females in the Con and BS exposed experimental groups. **F**. DML and DML-containing gene counts in F1 livers across the four sire groups and in males and females.

#### Sperm motility and epigenetics

Semen was collected from 16 sexually mature F1 rams (8 Con and 8 BS) representing the four sire family groupings (Table S4). Computer assisted sperm assessment found no effect of *in utero* BS exposure on either F1 (4.0 ± 0.53 vs 4.3 ± 0.36 billion/mL, Con vs BS) or F2 (2.3 ± 0.45 vs 1.5 ± 0.40 billion/mL, Con vs BS) sperm concentration. However, BS exposure increased (P = 0.044) the percentage of progressively motile F1 sperm (Fig. S2A) as well as F1 sperm straight-line velocity (86.7 ± 6.87 vs 105.9 ± 4.88 µm/sec, Con vs BS; P = 0.042).

For F1 sperm, RRBS reads covered 16,777,158 methylated cytosines. Hierarchical clustering of methylated cytosines revealed a strong sire influence within which was nested treatment (Fig. S3A). Subsequent analyses of DML split by sire revealed 47,921 unique DML, 80.5% of which were specific to a single sire. When averaged across sire lineages, 1,350 unique DML were observed, with a net decrease of 7.2% following BS exposure (Fig. S2B). Further analyses identified 150 genes containing genic DML, 12 of which have recognised roles in testis/sperm function (File S1C).

Micro RNAs (miRNAs) sequenced from seminal plasma and sperm were aligned to 153 mature sequences arising from the miRBase database prior to differential expression (DE) analyses (log_2_ fold change ≥ 2.0; FDR ≤ 0.01). When split by sire, analyses identified six unique miRNAs (miR-376e-5p, miR-376b-5p, miR-362, miR-544-5p, miR-539-5p and miR-376d). All six were DE in seminal plasma distributed across sire groups. miR-376b-5p was also DE in sperm.

### F2 epigenetic effects

#### Liver DNA methylation

DNA was analyzed from 15 male and 16 females (Table S5). RRBS covered 6,811,089 methylated cytosines. Hierarchical clustering of methylated cytosines revealed a stronger influence of sex, with treatment and grandsire effects nested within, than observed in F1 liver (Fig. S3C). When split by grandsire lineage 50,700 DML were observed, with 85.6% being specific to a single lineage (Fig. 2A), and with 1,794 DML (1,125 genic) common to those in F1 sperm. When split by sex, 1,256 DML were observed, with 98.2% specific to a single sex (Fig. 2B) *and* 13 DML (5 genic) common to those in F1 sperm. Mean methylation differences between treatments differed between sexes, with a 0.4% net decrease in males (Fig. 2C) and a 4.2% net increase in females (Fig. 2D) following BS exposure. Further analyses identified 174 unique genes containing genic DML, of which 48.3% were specific to male and 49.4% specific to female (File S1E, F).

**Figure 2.**
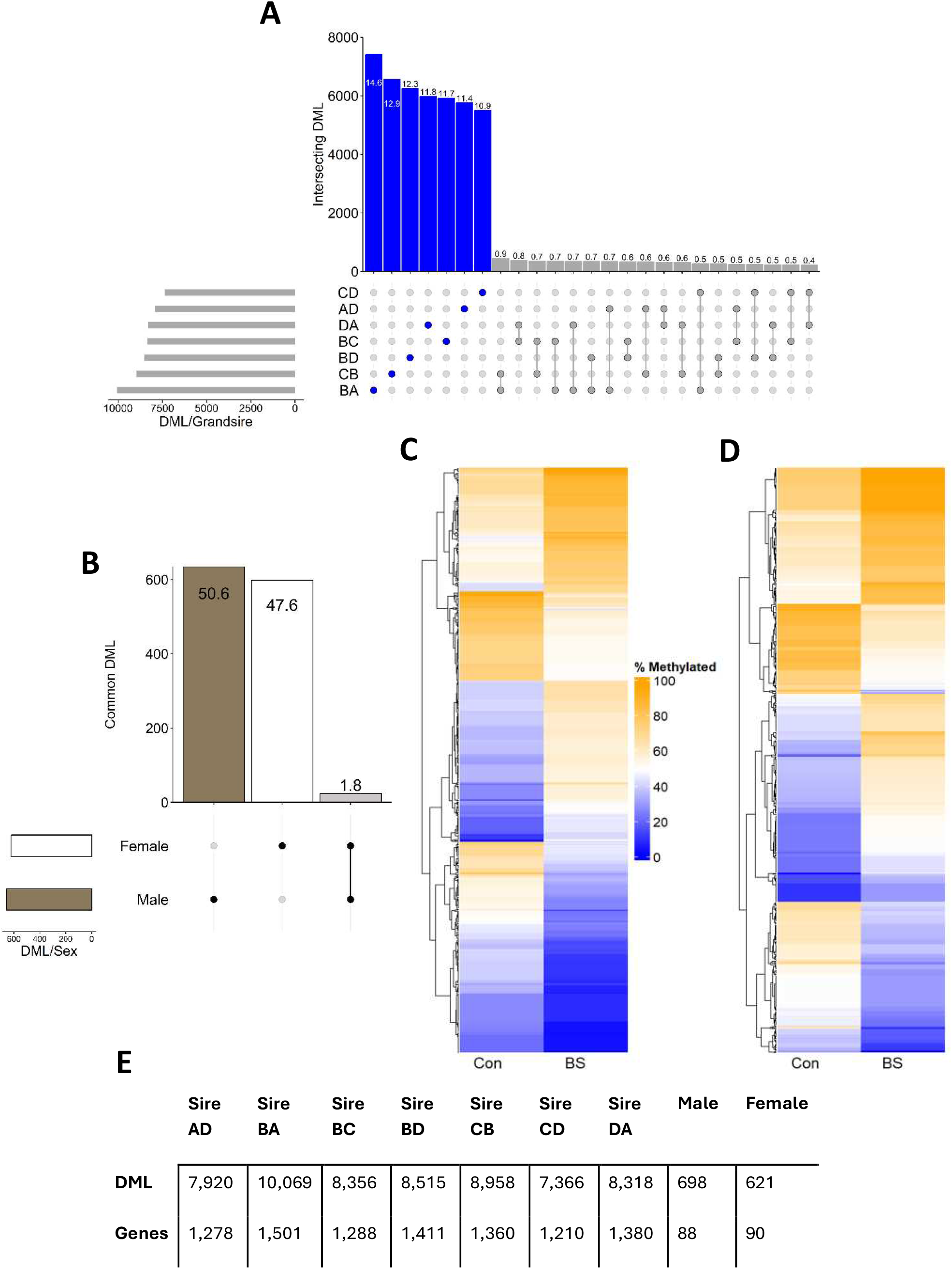
Methylation characteristics of F2 liver. **A.** Upset plot of overlapping differentially methylated loci (DML) (15% threshold) across seven F2 grandsire groups (Blue: sire specific, Grey: overlapping, values on bars denote percentage of total DML). Only the top 25 intersections are displayed. **B**. Upset plot of overlapping DML across male and female F2 liver (Black/White: sex specific, Grey: overlapping; values on bars denote percentage of total DML). **C**. Heatmap highlighting mean methylation of DML in males in the Control (Con) and Biosolids (BS) exposed experimental groups. **D**. Heatmap highlighting mean methylation of DML in females in the Con and BS exposed experimental groups. **E**. DML and DML-containing gene counts across seven F2 grandsire groups as well as in males and females.

#### Blood DNA methylation

DNA was also analyzed from 22 male and 22 female blood samples, representing almost all possible sire group combinations, balanced across the two treatments (Table S5). RRBS analyses covered 20,596,062 methylated cytosines. Hierarchical clustering revealed a similar pattern to F2 liver samples, showing treatment and grandsire lineage effects nested within strong sex effects (Fig. S3D). Differential methylation analyses identified 76,059 DML when split by grandsire, of which 71.7% were specific to a single grandsire (Fig. S4A). When split by sex 1,650 DML were observed of which 56.7% and 39.0% were unique to male and female respectively (Fig. S4B). Mean methylation differences between treatments, when averaged across grandsire lineage, differed between sexes, with a 10.1% net decrease in males (Fig. S4C) and a 3.0% net increase in females (Fig. S4D) following BS exposure. Further analyses of the 1,650 DML identified 181 unique genes containing genic DML, with 52.5% and 38.7% unique to male and female F2 offspring respectively (File S1G, H).

#### Sperm motility and epigenetics

Analyses were undertaken in semen from 14 sexually mature F2 rams (7 Con and 7 BS) representing seven matched grandsire family groupings (Table S5). In contrast to F1 sperm, there was no effect of BS exposure on percentage of progressive motility of F2 sperm (Fig. S2C) or on F2 sperm straight-line velocity (97.8 ± 14.85 vs 68.1 ± 13.29 µm/sec, Con vs BS). RRBS reads covered 14,876,916 methylated cytosines. Here, hierarchical clustering of methylated cytosines revealed a strong grand-paternal lineage influence within which was nested treatment (Fig. S3B). When averaged across grandsire lineages, 2,954 unique DMLs were observed, with a net increase of 2.2% following BS exposure (Fig. S2D). Further analyses identified 197 genes containing genic DML, 15 of which have recognised roles in testis/sperm function (File S1D). Finally, just under 2% of DML (n = 76) were common to both F1 and F2 sperm. Most were intergenic, but a cluster of 4 DML was located in Intron 3 of *GGH* and 11 DML in Intron 2 of *THRB*. There were also 12 genes harboring DML that were common to F1 and F2 sperm, one of which (*PTPRN2*) has a recognised role in testis/sperm function. In contrast to F1 sperm and seminal plasma, there were no DE miRNAs that reached the set criteria for FDR.

### F3 epigenetic effects

DNA was analyzed from 16 male and 16 female F3 offspring livers, balancing great-grandsire lineage across sex and treatment groups from the pool of available animals (Table S6). RRBS analyses covered 10,457,661 methylated cytosines. Hierarchical clustering again displayed strong separation by sex, with effects of treatment and great-grandsire lineage nested within sex (Fig. 3E). When split by great-grandsire lineage, 20,116 DML were identified, 89.5% of which were specific to a single great-grandsire lineage (Fig. 3A) with 187 DML (138 genic) common to those in F2 sperm. When split by sex, of 1,982 identified DML, 45.6% and 48.5% were specific to males and females respectively (Fig. 3B). Additionally, 67 of these DML (58 genic) were common to DML in F2 sperm. In males, there was a net methylation increase of 2.0% (Fig. 3C) whereas in females, there was a net methylation decrease of 7.1% following BS exposure (Fig. 3D). In addition, 362 unique genes were identified as containing genic DML, with 51.7% specific to males and 44.8% to females (File S1I, J). GO analyses of these 362 genes identified 11 enriched ‘biological processes’ terms in males (P ≤ 0.01) that encompass themes such as cell structure and adhesion, cell survival mechanisms and development and differentiation (Fig. S5A). Females had 35 enriched GO terms with common themes of cell adhesion and migration, development and morphogenesis and signaling (Fig. S5B).

**Figure 3.**
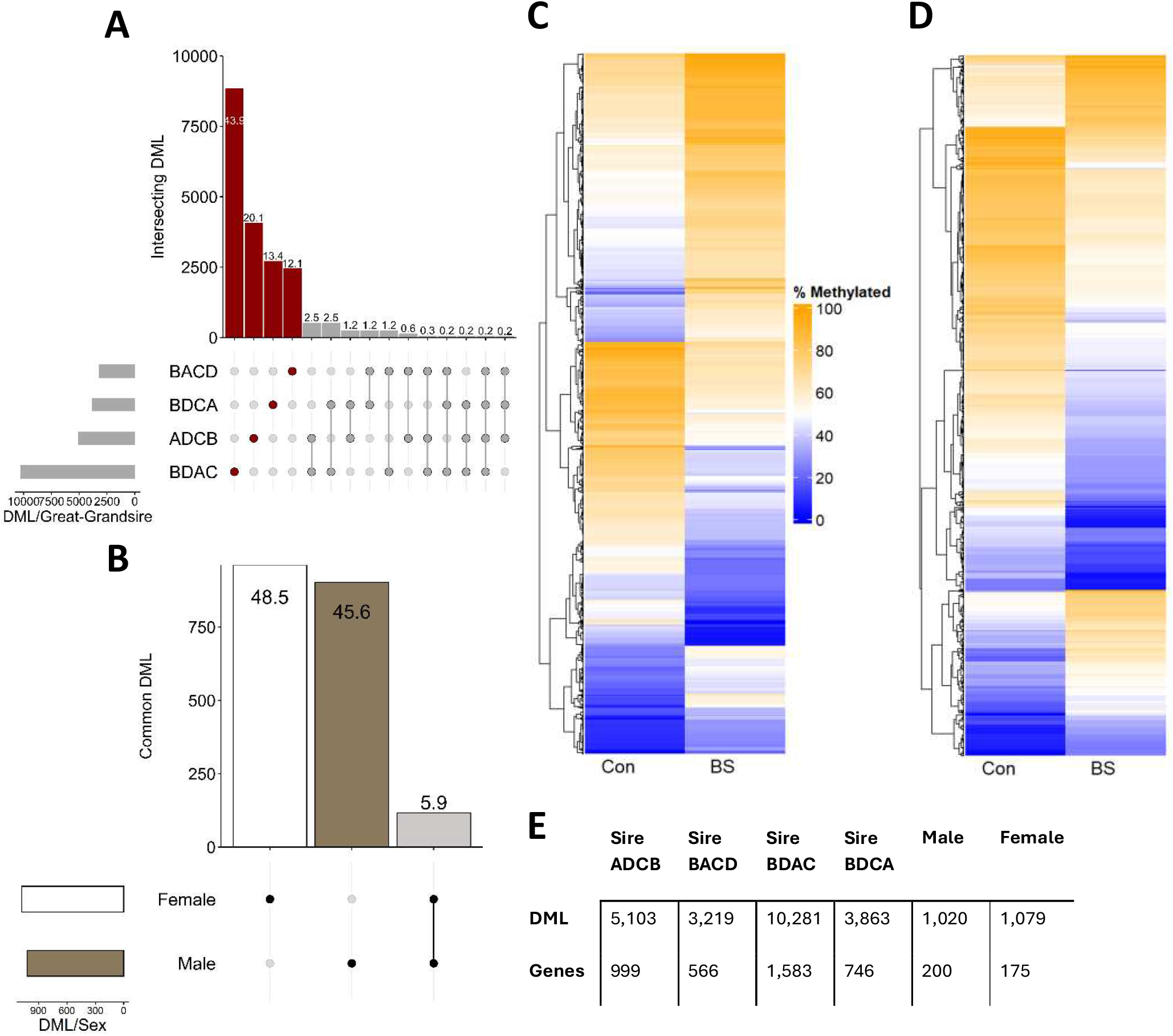
Methylation characteristics of F3 liver. **A.** Upset plot of overlapping differentially methylated loci (DML) (15% threshold) across four F3 great grandsire groups (Red: great grandsire specific, Grey: overlapping; values on bars denote percentage of total DML). **B**. Upset plot of overlapping DML across male and female F3 liver (Black/White: sex specific, Grey: overlapping; values on bars denote percentage of total DML). **C**. Heatmap highlighting mean methylation of DML in males in the Control (Con) and Biosolids (BS) exposed experimental groups. **D**. Heatmap highlighting mean methylation of DML (in females in the Con and BS exposed experimental groups. **E**. DML and DML-containing gene counts across four great-grandsire groups as well as in males and females.

### Differential methylation across generations

Initially, DML from each sex were compared across generations but no DML were found to be consistent across the three generations in either males (Fig. 4A) or females (Fig. 4B), other than a single genic DML region in a common and novel gene across the three generations in male (Fig. 4C) but not female (Fig. 4D) descendants.

**Figure 4.**
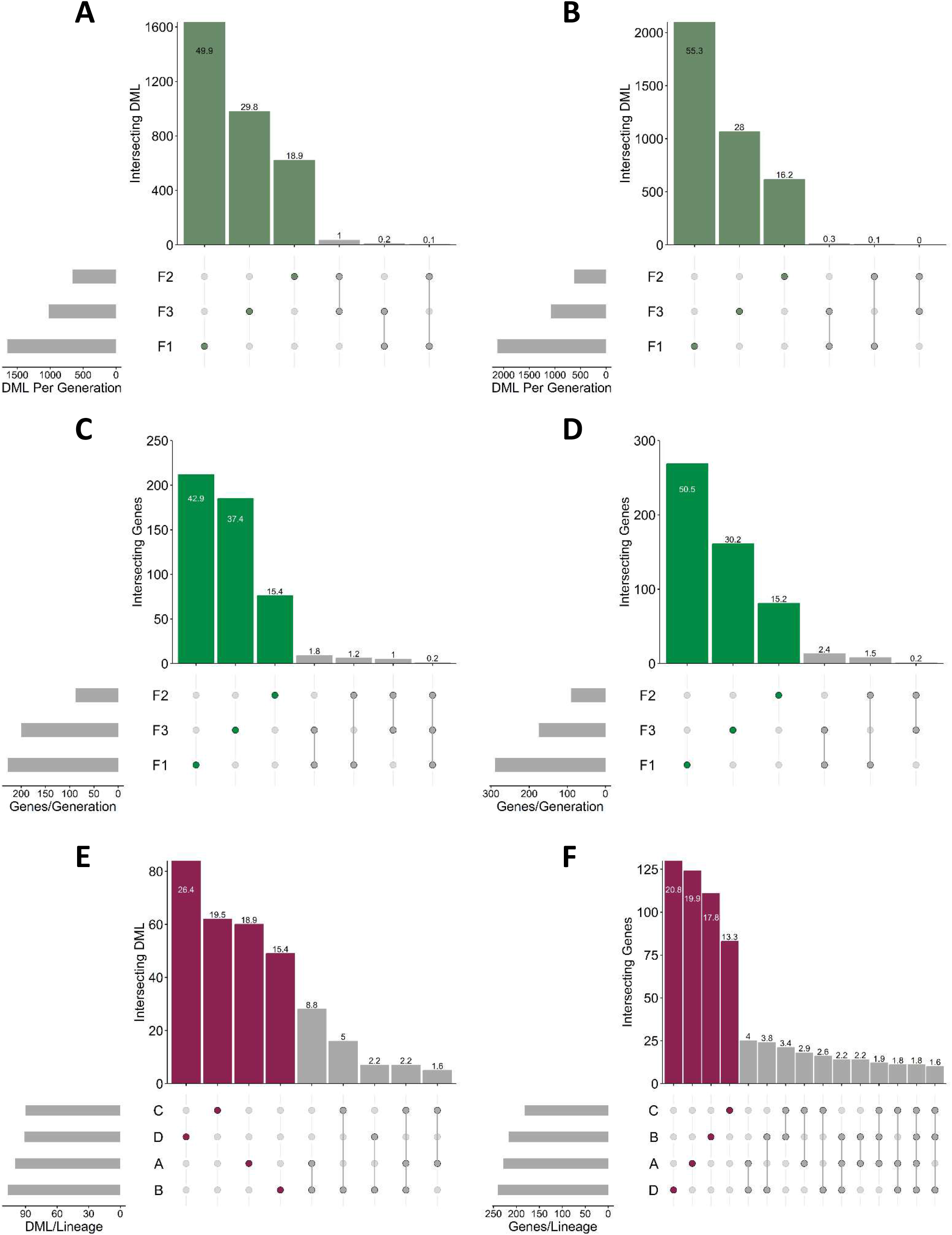
Intersecting differential methylated loci (DML) and genes containing DML in liver samples across generations. **A.** Intersecting DML across three generations of male offspring. **B**. Intersecting DML across three generations of female offspring. **C**. Intersecting DML-containing genes across three generations of male offspring. **D**. Intersecting DML-containing genes across three generations of female offspring. **E**. Intersecting cross-generational DML from each lineage (e.g., tracking all instances of F0 Sire A from F1, F2 and F3 analyses split by sire/grandsire/great-grandsire and averaged across sex), across the four lineages. **F**. Intersecting DML-containing genes present in all generations of a single lineage (e.g., tracking all instances of original Sire A from F1, F2 and F3 analyses split by sire/grandsire/great-grandsire and averaged across sex), across the four lineages.

An alternative approach split analyses by sire (F1), grandsire (F2), and great grandsire (F3) lineage, so that effects of sex within generation are averaged. This approach recognizes the significant proportion of DML that reside within descendants of each sire within each of the three generations (Fig. 1 to 3) and sought to track cross-generational DML within each founder-male lineage (i.e., descendants from F0 Rams A, B, C and D) to identify DML present in all 3 generations of each lineage. Grandsire and great-grandsire analyses were collated according to original F0 ram sire to produce single F2 and F3 DML lists (e.g., for lineage A: F1 = A; F2 = AD, BA, DA; F3 = ADCB, BACD, BDAC, BDCA) (Fig. S1). In total 100 DML across 16 genes were found in lineage A, 107 DML across 17 genes in lineage B, 90 DML across 15 genes in lineage C and 91 DML across 18 genes in lineage D. DML present in all three generations of a single lineage were then compared to those from other lineages to identify cross-generational DML that persisted regardless of lineage. None, however, were observed, although a 7 DML cluster within 39 bases within Intron 3 of *DHRSX* (just downstream of 3’UTR of *ZBED1;* Fig. S6A) was consistent for all three generations across lineages A, B and C (Fig. 4E). Adopting the same approach regarding genes containing DML, 11 genes were found to contain DML within all generations of all lineages. However, these DML were not consistent across generations within or across lineage. Instead, these genes (consisting of 6 known genes: *U6, MTCL1, PAQR5, ERG, SARDH, MGLL*)) represent common regions of the genome affected by differential methylation following BS exposure (Fig. 4F). Mapping DML for these 6 known genes revealed that, whilst they were generally clustered within each gene (e.g., DML within *PAQR5* were clustered within Intron 1), the exact position didn’t match on all occasions.

Finally, a third analysis separated male and female descendants. For each sex, all samples from all three generations were analyzed. This involved averaging effects of sire, grandsire, and great-grandsire lineage (as presented in Fig. 4E and 4F), as well as effects arising from each generation. This identified 97 cross-generational DML in males (Fig. S7A) and 19 in females (Fig. S7B). DML positions in males and females were then compared between sex (Fig. 7C), showcasing absolute sexual-dimorphism in cross-generational DML. These analyses identified 9 known genes (including *DHRSX*) and 4 novel genes in males. DML clusters were observed in four of these genes (i.e., *PGRMC1, 5SrRNA, ME3/SNORD30* and *DHRSX*). For *DHRSX*, 8 DMLs clustered in Intron 3, seven of which matched those identified cross-generationally in Lineages A, B and C (Fig. 4E). These analyses also identified two known (*DPP6* and *COL9A1*) and three novel genes in females. As *DHRSX* was identified in two separate analyses, transcript expression for this gene was determined in F3 liver. *DHRSX* expression was increased (P < 0.01) in F3 male (Fig. S7D) but not F3 female (Fig. S7E) descendants following *in utero* exposure of the F1 generation to chemicals originating from BS. This was consistent with a 28% decrease in methylation for all 7 DML in Intron 3 in F3 males with no change in methylation at the corresponding locus in females.

### ‘Escapees’

High-confidence demethylation-resistant genes within the germline (termed ‘escapees’) have been identified in the pig (31), a proportion of which have conserved synteny with both humans and mice (32,33). From the published list of 101 escapees, 93 named candidates were cross-referenced to the list of hepatic F3 DML.

Initial analysis split sex and averaged between great-grandsire lineages. Relative to regular genes, the Odds Ratio of ‘escapees’ containing a DML was greater than 5.1 (P < 0.01). However, only a few F3 DML within ‘escapees’ were identified per sex, with none consistently represented across all three generations. Next, an analysis using DML obtained by averaging across sex and splitting by great grandsire lineage identified 219 DML in 30 ‘escapees’ (Fig. S8A). In this case, the Odds Ratio of ‘escapees’ containing a DML was significant (P < 0.001) but varied between great-grandsire lineage (ranging from 2.9 to 4.3). Then, averaging across sex, but including F1 and F2 generations along with F3, revealed 20 ‘escapees’ (including *DPP6*) that contained DML identifiable across all three generations (Fig. S8C; File S1K). However, these DML were not necessarily the same in all three generations although, importantly, further analysis did identify 10 DML that persisted across all three generations (Fig. S8B). Intriguingly, these DML all clustered within 169 bases of intron 4 of the tumor suppressor gene *CADM1* (Fig. S6B) with one DML in a region showing conservation with a human *CADM1* CTCF binding site involved in transcriptional regulation (34). Looking back at the F3 analysis split by sex and averaged across great grandsire, these 10 DML were not detectable on average in either sex at the 15% threshold, but present across F3 males only with a mean 9% increase in methylation. Importantly, transcript expression for *CADM1* was reduced (P = 0.043) in F3 male (Fig. S8D) but not F3 female (Fig. S8E) descendants.

Finally, germline cross-generational analyses of overlapping DML between F1 sperm and F2 liver and blood, and F2 sperm and F3 liver, for the listed ‘escapees’ revealed 32 DML in 5 ‘escapees’ in F1 sperm (Fig. S9A). In this instance, the two DML observed in sperm *DPP6* were also identified in both F2 liver and blood. These analyses also revealed 80 DML in 5 ‘escapees’ in F2 sperm (Fig. S9B). Here, nine DML in sperm *DPP6*, and 6 DML in sperm *COL5A1*, were also identified in F3 liver.

## Discussion

Several important and novel findings emerge from the current study. First, we demonstrate that gestational exposure to a real-life mixture of EC, using an established outbred animal model of the human exposome (1), leads to measurable alterations to DNA methylation in F1 offspring, with detectable effects in F2 and F3 descendants. Equally, our data reveal the sizable extent to which genetic ancestry and offspring sex influence DNA methylation. Depending on tissue type, between 70 - 94% of DML within each generation were specific to a single sire lineage, and 88 - 96% of DML were sex specific. Such strong lineage and sex effects complicate interpretation and highlight the difficulty of distinguishing EC exposure-related epigenetic change from genetically regulated methylation variation (24,25). The progressive convergence of F0 paternal lineage across generations (Fig. S1) was reflected in cytosine methylation clustering (Fig. 2), with sex and treatment effects becoming more pronounced in later generations. A key challenge is determining whether observed effects on DNA methylation constitute transgenerational epigenetic inheritance (TEI). In part, this requires (i) propagation of EC induced epigenetic marks through the germline beyond the directly exposed generations, (ii) recurrence of the same EC induced epimutations in the first unexposed generation (F3 in this study), and (iii) associated differences in gene expression (35). Based on these criteria alone, our data do not provide definitive evidence for TEI, although they highlight regions where environmentally associated methylation differences appear repeatedly across generations. Further analyses would be required to link observed DNA methylation differences in F1 and F2 sperm, and miRNA in F1 sperm and seminal plasma, with altered methylation in subsequent somatic cell lineages. Furthermore, although persistently altered methylation was identified at specific loci, such as regions within *DHRSX* and *CADM1* (both of which were differentially expressed in F3 descendants), these changes were not uniformly present across all lineages. Such inconsistency, combined with RRBS’s inherent genomic bias and the strong influence of allele-specific methylation (ASM) and methylation quantitative trait loci (meQTLs) effects (36,37), limits our ability to attribute these recurrent loci to stable transgenerational transmission rather than genetically driven variation. Nevertheless, these findings illustrate that complex, low-dose EC mixtures can induce robust methylation changes that recur in multiple generations, in an outbred mammalian species, and point to specific genomic regions where exposure-related perturbations arise repeatedly across generations, and which warrant targeted investigation in future TEI-focused studies.

### DNA methylation sequencing in an outbred species

A key consideration in interpreting our findings is the sequencing approach used. RRBS enriches CpG-dense regions, that include CpG islands, promoter-proximal areas, and 5’ regulatory elements, via *MspI* digestion at 5′-C^CGG-3′ sites, which are over-represented in these genomic contexts (38). In our study, this was reflected by the consistent distribution of hepatic DML between islands, shores, shelves and open sea (43.0%, 11.0%, 5.2% and 40.5%, respectively). RRBS therefore provides a non-random and promotor biased view, with limited coverage of distal enhancers and repetitive elements, regions that may be sensitive to environmental exposures (39). Instead, RRBS preferentially captures stable, cis-regulated CpGs and promoter-related methylation changes, which often reflect durable early-life programming effects (40) and have been implicated previously in developmental responses to toxicants in fish and mice (41-44). For the scale and given constraints of the current study, RRBS facilitated greater sequencing depth and biological replication across multiple generations and different cell types than would have otherwise been possible using alternative genome-wide single-nucleotide resolution platforms (45). Adequate depth and replication are essential in outbred species, where a substantial proportion of methylation variation is genetically influenced, including through ASM and meQTLs (25,36,37). Our controlled breeding structure, which matched F0 sire lineages within and across generations, helped mitigate, but cannot eliminate, these confounding effects (46,47). Filtering CpGs overlapping known SNPs further reduced artefactual differential methylation arising from CpG-SNPs (48). However, the effectiveness of this filtering depends on the completeness of available SNP catalogues and local linkage disequilibrium. Un-assayed causal variants in proximity to measured CpGs could still generate meQTL-like differences even when the CpG itself is invariant.

### Sexual dimorphism in DNA methylation inheritance

Several studies have reported sexually dimorphic DNA methylation responses following *in utero* and perinatal exposure to EC and heavy metals using RRBS in rodent offspring liver (43,49,50) and heart (51). In our study, the magnitude of sex-specific differences in hepatic DNA methylation across generations following *in utero* EC exposure was comparable to the influence of sire lineage itself. Notably, there is no evidence that RRBS inherently exaggerates sex differences compared with other single-nucleotide-resolution platforms (52). As RRBS captures CpGs mainly within promoter-proximal and CpG-island regions, while incompletely sampling distal enhancers, intronic elements, and open-sea regulatory sites (53), our findings should be interpreted primarily within this regulatory context. Many hepatic genes with stable, sex-biased methylation patterns in mammals, including those regulating lipid metabolism, energy homeostasis, and endocrine pathways (e.g., *IGF1, NR3C1, PPARA*), are influenced by methylation in these promoter-proximal regions (54-56). Although these canonical loci did not feature in our cross-generational gene sets, 68 other metabolic genes did (File S1). Most were male-biased, with only nine expressed in females. Among these, *IMMP1L, TFPI*, and *DPP6* contribute to hepatic proteolysis and nitrogen metabolism, while *GALNT17, DEPTOR, PDZRN3*, and *EEFSEC* regulate transcription, translation, or protein synthesis. Functional enrichment further highlighted divergence between sexes: male-biased genes were enriched for processes such as cell adhesion, apoptosis, oxidative phosphorylation, and carbohydrate utilisation; female-biased genes were more strongly associated with developmental signalling pathways (e.g., NOTCH, WNT, BMP), lipid differentiation, and peroxisomal dynamics. Both sexes shared enrichment for homophilic cell adhesion, suggesting a conserved biological theme achieved through sex-specific molecular networks. These sexually dimorphic effects could have arisen because of sex-chromosome-directed epigenetic alterations to autosomal DNA methylation and/or due to the actions of sex steroids (26). Given that EC exposure extended throughout gestation it is not possible at this juncture to ascertain which of these mechanisms may have been more prominent.

### Multi-generational epigenetic inheritance

*In utero* exposure to endocrine disruptors and other toxicants has altered the profile of small non-coding RNA (sncRNA) of adult F1 sperm in laboratory species (57-59). Furthermore, semen-borne miRNA can contribute to paternal epigenetic signalling, consistent with recognised modes of intergenerational inheritance, although mechanistic details remain uncertain (60). Notably, while gestational toxicant exposure can induce sperm miRNA differences that persist to F3 in outbred rats (61), the six annotated sheep miRNAs detected here were restricted to F1 semen. One of these, miR-539-5p, is known to regulate DNA methylation in mice by targeting *Dnmt3b*, thereby modifying promoter CpG island methylation and transcription of lipogenic genes such as *Srebf1* (62). Whilst no comparable role has been defined for the remaining five microRNA, miR-544-5p forms part of a cluster of miRNA within the DLK1-GTL2/MEG3 imprinted domain on sheep chromosome 18, with miR-376b-5p, miR-376e-5p and miR-376d residing close by (63). This domain regulates the callipyge muscle hypertrophy phenotype in sheep (64,65). However, no differential methylation at this locus was detected in our datasets (File S1), consistent with its muscle-specific regulation (65).

The current study identified a number of overlapping DML between F1 sperm and F2 liver or blood, and between F2 sperm and F3 liver. However, care is required when ascribing these as inherited differences. The current dataset cannot distinguish true inheritance from coincidental coverage or meQTL-driven methylation. Of the genes listed as harboring overlapping DML (File S1), however, *SNED1* (with 45 overlapping DML) and *DPP6* (a listed ‘‘escapee’ with 9 overlapping DML) are worthy of further investigation. Also, *PTPRN2* (Fig. S8A), another listed ‘escapee’ that overlapped between F1 and F2 sperm. Although it did not feature in the list of DE genes for the testis in the current cohort of F1 animals (12), hypermethylation and reduced expression of this gene in sperm is associated with reduced sperm counts and motility in men exposed to high levels cigarette chemicals (66,67).

When broader functional sets were examined, pairwise comparisons of differentially methylated hepatic genes (File S1) across generations revealed consistent enrichment for ‘cell organisation’, ‘cell adhesion’, and ‘lipid/redox metabolism’ in males. Intriguingly, plasma and liver triglyceride concentrations were both increased in males, but not females, among the broader cohort of our participating F1 offspring (21). In females, overlapping gene sets were enriched for ‘cell adhesion’, ‘neurodevelopmental signalling’, and ‘transcriptional regulation’. Although underlying DML and gene identities varied between generations, the stability of these functional themes suggests that multigenerational liver responses may be conserved at the process level rather than through persistence of identical epimutations.

A key criterion for TEI relates to the inheritance of the same epimutation(s) across generations extending to the first unexposed generation (F3 in this study) (35). Two genes from our dataset warrant consideration in this respect. The ‘escapee’ *CADM1* harbored 10 DML clustered in a conserved intronic region of the gene which, in humans, is associated with transcriptional regulation (34). This cluster was detected in at least one lineage across all three generations (Fig. S6B), and increased methylation in F3 males was associated with reduced transcription (Fig. S8D). *CADM1* is a recognized tumor suppressor gene regulating hepatic cell-cycle progression (68) and mediating endothelial adhesion and liver inflammation (69). Interestingly, prenatal testosterone exposure in adult female sheep altered methylation at a separate intronic DMR of *CADM1* (located in Intron 2) but with no effect on transcription (70), supporting its sensitivity to developmental endocrine perturbation. The second gene, *DHRSX*, is involved in the dolichol pathway, linked to glycosylation and hepatic glycoprotein production (71). Importantly, it is located in the pseudo-autosomal region of the X-chromosome in both humans and sheep (72,73) and thus escapes inactivation. We identified 7 DML clustered within Intron 3 (Fig. S6A) that recurred in all three generations in lineages A, B and C (Fig. 4E), although the underlying cause cannot be distinguished. Furthermore, reduced methylation at this locus in F3 males aligned with increased transcript expression (Fig. S7D). This methylation pattern was consistent across all generations of males in lineages B and C, but not in lineage A, suggesting partial genetic-inherited regulation of methylation at this site. A putative imprinting control region located in human *DHRSX* Intron 3 (74), which appears orthologous to the sheep locus, raises the possibility of parent-of-origin effects contributing to the observed sexual dimorphism.

### Concluding remarks

This study demonstrates that gestational exposure to low-level complex mixtures of EC, representative of the human exposome, can induce sexually dimorphic and lineage-specific changes in DNA methylation that can recur across multiple generations in an outbred large-animal model. Although no single differentially methylated locus persisted consistently in all lineages across all generations, the identification of recurrent changes in regions within defined genes such as *DHRSX* and *CADM1* provides intriguing, though not definitive, evidence for multi-generational recurrence of exposure-induced epimutations. These findings underscore the challenge of disentangling true transgenerational DNA methylation inheritance from underlying genetic lineage-directed variation. Missing from the current dataset are metabolic and reproductive phenotype data linked to DML for F2 and F3 descendants. These are required to demonstrate transgenerational inheritance (35) and are the subject of ongoing investigations with the current cohort of animals. Conceptually, however, findings from our study concur with the conclusion reached, following a series of TEI studies in sheep, that genomic context is a key determinant of how methylation changes persist across generations following initial environmental exposure (75-77). Consequently, future research will expand our initial observations to whole-genome approaches, to capture distal enhancer and repeat sequences. Germline transmission analyses will be expanded to include oocytes, and to assess chromatin accessibility using contemporary single cell multiomic approaches. Dense genotyping combined with whole genome sequencing will map meQTL and ASM, testing methylation using pedigree-aware mixed models to partition genetic and exposure effects. Ultimately, however, further reciprocal matings across lineages may be required to disentangle sire/grandsire contributions and assess DML persistence within haplotypes.

## Materials and Methods

Details can be found in SI Appendix, Detailed Materials and Methods. Animal procedures were approved by the Animal Welfare and Ethical Review Board of the University of Glasgow and performed under the United Kingdom’s Animals (Scientific Procedures) Act 1986 (licensed authority: PF10145DF). Protocols complied with the ARRIVE guidelines.

### Experimental treatments, animals and design

EasyCare ewes (*Ovis aries*; F0, n = 320) were allocated randomly to graze either Con or BS treated pastures at the University of Glasgow Cochno Farm and Research Centre (7) from six weeks prior to mating (by artificial insemination; AI) until lambing, then grazed on Con pastures thereafter. AI (78) was undertaken in 307 F0 ewes using semen from four unrelated F0 rams (labelled A, B, C and D) not exposed to BS, generating four F1 family groups matched across treatments (Fig. S1, Table S1) (15). Subsequently, semen from eight F1 rams (4 Con and 4 BS), from each of the four families within treatment group, was used for AI of 149 F1 ewes from the other three families (Fig. S1, Table S2). Natural matings between F2 rams and ewes (n = 61) were undertaken within each treatment group to generate F3 offspring (Fig. S1, Table S3).

### Computer assisted sperm assessment (CASA) and semen cryopreservation

Semen was collected from 16 F1 and 14 F2 sexually mature rams on two occasions within generation, balancing sire lineage between treatments (Table S1). Sperm analyses were undertaken using iSperm mCASA (Aidmics Biotechnology Co., Ltd, Taipei, Taiwan) (79). Residual seminal plasma and sperm were stored at -80°C.

### miRNA analyses from sperm and seminal plasma

Following extraction (miRNeasy kit; Qiagen) with DNase treatment (80) sperm RNA libraries were pooled and sequenced on the Element Biosciences AVITI System on an AVITI 2X75 Sequencing Kit - Cloudbreak FS Medium Output (Element Biosciences; 860-00014), generating approximately 20 million 75-bp single-end reads per sample. Raw sequencing reads were aligned to mature and precursor sheep miRNA sequences from the miRBase database (81). Differential expression (DE) performed on miRNAs with > 10 counts per million using the Bioconductor edgeR package v4.0.16 (82). DE miRNAs = log fold change ≥ 2.0, FDR ≤ 0.01 (83).

### DNA extraction

DNA was extracted (DNeasy Blood and Tissue Kit (Qiagen) with RNase A (for liver/blood), and gradient plus somatic cell lysis (with DNase) followed by salting-out (84) (for sperm).

### Reduced Representation Bisulfite Sequencing

DNA and unmethylated λ (Promega, Southampton, UK) were digested with *Msp1* (NEB, Hitchin, uk). Methylated adaptor was ligated to DNA. RRBS libraries were then prepared using bisulfite-converted DNA. Size-selected fragments (200-600 bp) were pooled and sequenced (Illumina NovoSeq X plus), > 80 million 150 bp pair-end reads per sample (Novogene, Cambridge, UK).

### Differential Methylation Analyses

Sequenced RRBS libraries were aligned to the Ensembl reference genome ARS-UI_Ramb_v2.0 (GCA_016772045.1). CpGs matching positions of known SNPs were filtered. Analyses were performed using DSS (85) with a threshold ≥ 15% difference in methylation, FDR ≤ 0.01 (83). Identified DMLs labelled genic or intergenic using the ChIPseeker v1.34.1 package (86,87). GO and KEGG were performed on genic DMLs. ‘Escapees’ (31) were identified and enrichment determined using Fisher’s exact test.

### Statistical analysis

Used GenStat statistical package (21^st^ Edition, VSN International, 2022; https://www.vsni.co.uk/). Sperm motility, transcript expression (F3 livers) analyzed using REML generalized linear mixed models (fixed term = treatment; random terms = lineage group (F1, F2 or F3) Data are presented as predicted means ± SEM.

## Acknowledgments

The graphical figure (Fig. S1) was created with BioRender.com.

## Data Availability

All study data are included in the article, SI Appendix, and Data File S1. RRBS and RNA sequencing data have been deposited in the European Nucleotide Archive and are accessible through accession number PRJEB109928.

## References

1. M. Bellingham, N. P. Evans, R. G. Lea, V. Padmanabhan, K. D. Sinclair, Reproductive and Metabolic Health Following Exposure to Environmental Chemicals: Mechanistic Insights from Mammalian Models. Annu Rev Anim Biosci 13, 411–440 (2025).

2. N. Güil-Oumrait et al., Prenatal Exposure to Chemical Mixtures and Metabolic Syndrome Risk in Children. JAMA Netw Open 7, e2412040 (2024).

3. N. Robles-Matos, T. Artis, R. A. Simmons, M. S. Bartolomei, Environmental Exposure to Endocrine Disrupting Chemicals Influences Genomic Imprinting, Growth, and Metabolism. Genes (Basel) 12 (2021).

4. E. E. Nilsson, M. Ben Maamar, M. K. Skinner, Role of epigenetic transgenerational inheritance in generational toxicology. Environ Epigenet 8, dvac001 (2022).

5. B. Sharma, A. Sarkar, P. Singh, R. P. Singh, Agricultural utilization of biosolids: A review on potential effects on soil and plant grown. Waste Manag 64, 117–132 (2017).

6. M. Bellingham, N. Evans, Impact of Real-Life Environmental Exposures on Reproduction: Biosolids and Male Reproduction. Reproduction 168 (2024).

7. M. Bellingham et al., Exposure to a complex cocktail of environmental endocrine-disrupting compounds disturbs the kisspeptin/GPR54 system in ovine hypothalamus and pituitary gland. Environ Health Perspect 117, 1556–1562 (2009).

8. M. Bellingham et al., Exposure to chemical cocktails before or after conception---the effect of timing on ovarian development. Mol Cell Endocrinol 376, 156–172 (2013).

9. R. G. Lea et al., The fetal ovary exhibits temporal sensitivity to a ‘real-life’ mixture of environmental chemicals. Sci Rep 6, 22279 (2016).

10. R. G. Lea et al., Ovine fetal testis stage-specific sensitivity to environmental chemical mixtures. Reproduction 163, 119–131 (2022).

11. M. Bellingham et al., Foetal and post-natal exposure of sheep to sewage sludge chemicals disrupts sperm production in adulthood in a subset of animals. Int J Androl 35, 317–329 (2012).

12. C. S. Elcombe et al., Developmental exposure to real-life environmental chemical mixture programs a testicular dysgenesis syndrome-like phenotype in prepubertal lambs. Environ Toxicol Pharmacol 94, 103913 (2022).

13. C. S. Elcombe et al., Developmental exposure to a real-life environmental chemical mixture alters testicular transcription factor expression in neonatal and pre-pubertal rams, with morphological changes persisting into adulthood. Environ Toxicol Pharmacol 100, 104152 (2023).

14. M. Ghasemzadeh Hasankolaei et al., In-utero exposure to real-life environmental chemicals disrupts gene expression within the hypothalamo-pituitary-gonadal axis of prepubertal and adult rams. Environ Res 264, 120303 (2025).

15. N. P. Evans et al., Sexually dimorphic impact of preconceptional and gestational exposure to a real-life environmental chemical mixture (biosolids) on offspring growth dynamics and puberty in sheep. Environ Toxicol Pharmacol 102, 104257 (2023).

16. K. M. Halloran et al., Developmental programming: preconceptional and gestational exposure of sheep to biosolids on offspring ovarian dynamics†. Biol Reprod 112, 331–345 (2025).

17. Y. Zhou et al., Developmental programming: mechanisms of early exposure to real-life chemicals in biosolids on offspring ovarian dynamics†. Biol Reprod 112, 1229–1242 (2025).

18. S. V. Thangaraj et al., Developmental programming: Preconceptional and gestational exposure of sheep to a real-life environmental chemical mixture alters maternal metabolome in a fetal sex-specific manner. Sci Total Environ 864, 161054 (2023).

19. S.V. Thangaraj et al., Developmental programming: Impact of preconceptional and gestational exposure to a real-life environmental chemical mixture on maternal steroid, cytokine and oxidative stress milieus in sheep. Sci Total Environ 900:165674. (2023).

20. M. Ghasemzadeh-Hasankolaei et al., Preconceptional and in utero exposure of sheep to a real-life environmental chemical mixture disrupts key markers of energy metabolism in male offspring. J Neuroendocrinol 36, e13358 (2024).

21. S. V. Thangaraj et al., Developmental programming: Sex-specific effects of prenatal exposure to a real-life mixture of environmental chemicals on liver function and transcriptome in sheep. Environ Pollut 367, 125630 (2025).

22. R. P. Thompson, E. Nilsson, M. K. Skinner, Environmental epigenetics and epigenetic inheritance in domestic farm animals. Anim Reprod Sci 220, 106316 (2020).

23. O. Van Cauwenbergh, A. Di Serafino, J. Tytgat, A. Soubry, Transgenerational epigenetic effects from male exposure to endocrine-disrupting compounds: a systematic review on research in mammals. Clin Epigenetics 12, 65 (2020).

24. N. E. Banovich et al., Methylation QTLs are associated with coordinated changes in transcription factor binding, histone modifications, and gene expression levels. PLoS Genet 10, e1004663 (2014).

25. S. Villicaña, J. T. Bell, Genetic impacts on DNA methylation: research findings and future perspectives. Genome Biol 22, 127 (2021).

26. K. D. Sinclair, International Symposium on Ruminant Physiology: Developmental epigenetics-Understanding genetic and sexually dimorphic responses to parental diet and outcomes following assisted reproduction. J Dairy Sci 108, 7723–7740 (2025).

27. M. H. Fitz-James, G. Cavalli, Molecular mechanisms of transgenerational epigenetic inheritance. Nat Rev Genet 23, 325–341 (2022).

28. A. J. Drake et al., In utero exposure to cigarette chemicals induces sex-specific disruption of one-carbon metabolism and DNA methylation in the human fetal liver. BMC Med 13, 18 (2015).

29. C. McCabe, O. S. Anderson, L. Montrose, K. Neier, D. C. Dolinoy, Sexually Dimorphic Effects of Early-Life Exposures to Endocrine Disruptors: Sex-Specific Epigenetic Reprogramming as a Potential Mechanism. Curr Environ Health Rep 4, 426–438 (2017).

30. Q. Chen et al. Sperm tsRNAs contribute to intergenerational inheritance of an acquired metabolic disorder. Science 351, 397–400 (2016).

31. Q. Zhu et al., Specification and epigenomic resetting of the pig germline exhibit conservation with the human lineage. Cell Rep 34, 108735 (2021).

32. W. W. Tang et al., A Unique Gene Regulatory Network Resets the Human Germline Epigenome for Development. Cell 161, 1453–1467 (2015).

33. S. Guibert, T. Forné, M. Weber, Global profiling of DNA methylation erasure in mouse primordial germ cells. Genome Res 22, 633–641 (2012).

34. K. Campos-León et al., Repression of CADM1 transcription by HPV type 18 is mediated by three-dimensional rearrangement of promoter-enhancer interactions. PLoS Pathog 21, e1012506 (2025).

35. H. Khatib, J. Townsend, M. A. Konkel, G. Conidi, J. A. Hasselkus, Calling the question: what is mammalian transgenerational epigenetic inheritance? Epigenetics 19, 2333586 (2024).

36. C. Do et al., Mechanisms and Disease Associations of Haplotype-Dependent Allele-Specific DNA Methylation. Am J Hum Genet 98, 934–955 (2016).

37. H. Schulz et al., Genome-wide mapping of genetic determinants influencing DNA methylation and gene expression in human hippocampus. Nat Commun 8, 1511 (2017).

38. A. Meissner et al., Genome-scale DNA methylation maps of pluripotent and differentiated cells. Nature 454, 766–770 (2008).

39. S. J. Shareef et al., Extended-representation bisulfite sequencing of gene regulatory elements in multiplexed samples and single cells. Nat Biotechnol 39, 1086–1094 (2021).

40. T. Hachiya et al., Genome-wide identification of inter-individually variable DNA methylation sites improves the efficacy of epigenetic association studies. NPJ Genom Med 2, 11 (2017).

41. A. Brulport, D. Vaiman, M. C. Chagnon, L. Le Corre, Obesogen effect of bisphenol S alters mRNA expression and DNA methylation profiling in male mouse liver. Chemosphere 241, 125092 (2020).

42. K. Nohara et al., Gestational arsenic exposure induces site-specific DNA hypomethylation in active retrotransposon subfamilies in offspring sperm in mice. Epigenetics Chromatin 13, 53 (2020).

43. K. Wang et al., Tissue-and Sex-Specific DNA Methylation Changes in Mice Perinatally Exposed to Lead (Pb). Front Genet 11, 840 (2020).

44. A. S. Voisin, V. Suarez Ulloa, P. Stockwell, A. Chatterjee, F. Silvestre, Genome-wide DNA methylation of the liver reveals delayed effects of early-life exposure to 17-α-ethinylestradiol in the self-fertilizing mangrove rivulus. Epigenetics 17, 473–497 (2022).

45. R. Doherty, C. Couldrey, Exploring genome wide bisulfite sequencing for DNA methylation analysis in livestock: a technical assessment. Front Genet 5, 126 (2014).

46. K. B. Michels et al., Recommendations for the design and analysis of epigenome-wide association studies. Nat Methods 10, 949–955 (2013).

47. A. F. McRae et al., Identification of 55,000 Replicated DNA Methylation QTL. Sci Rep 8, 17605 (2018).

48. M. B. C. Maldonado et al., Identification of bovine CpG SNPs as potential targets for epigenetic regulation via DNA methylation. PLoS One 14, e0222329 (2019).

49. Liu S. et al. “Perinatal DEHP exposure induces sex-and tissue-specific DNA methylation changes in both juvenile and adult mice.” Environmental Epigenetics 7(1): dvab004 (2021).

50. D. L. Maxwell et al., Mixtures of per-and polyfluoroalkyl substances (PFAS) alter sperm methylation and long-term reprogramming of offspring liver and fat transcriptome. Environ Int 186, 108577 (2024).

51. L. K. Svoboda et al., Perinatal Lead Exposure Promotes Sex-Specific Epigenetic Programming of Disease-Relevant Pathways in Mouse Heart. Toxics 11 (2023).

52. X. Liu et al., Beyond the base pairs: comparative genome-wide DNA methylation profiling across sequencing technologies. Brief Bioinform 25 (2024).

53. K. Wreczycka et al., Strategies for analyzing bisulfite sequencing data. J Biotechnol 261, 105–115 (2017).

54. K. A. Lillycrop, E. S. Phillips, A. A. Jackson, M. A. Hanson, G. C. Burdge, Dietary protein restriction of pregnant rats induces and folic acid supplementation prevents epigenetic modification of hepatic gene expression in the offspring. J Nutr 135, 1382–1386 (2005).

55. Q. Fu, X. Yu, C. W. Callaway, R. H. Lane, R. A. McKnight, Epigenetics: intrauterine growth retardation (IUGR) modifies the histone code along the rat hepatic IGF-1 gene. Faseb j 23, 2438–2449 (2009).

56. D. J. Carr, J. S. Milne, R. P. Aitken, C. L. Adam, J. M. Wallace, Hepatic IGF1 DNA methylation is influenced by gender but not by intrauterine growth restriction in the young lamb. J Dev Orig Health Dis 6, 558–572 (2015).

57. M. Ben Maamar et al., Alterations in sperm DNA methylation, non-coding RNA expression, and histone retention mediate vinclozolin-induced epigenetic transgenerational inheritance of disease. Environ Epigenet 4, dvy010 (2018).

58. P. M. Herst et al., Folic acid supplementation reduces multigenerational sperm miRNA perturbation induced by in utero environmental contaminant exposure. Environ Epigenet 5, dvz024 (2019).

59. H. Hamada et al., Prenatal maternal glucocorticoid exposure modifies sperm miRNA profiles across multiple generations in the guinea-pig. J Physiol 602, 2127–2139 (2024).

60. A. Lismer, S. Kimmins, Emerging evidence that the mammalian sperm epigenome serves as a template for embryo development. Nat Commun 14, 2142 (2023).

61. H. McSwiggin, R. Magalhães, E. E. Nilsson, W. Yan, M. K. Skinner, Epigenetic transgenerational inheritance of toxicant exposure-specific non-coding RNA in sperm. Environ Epigenet 10, dvae014 (2024).

62. D. Kesharwani, A. Kumar, A. Rizvi, M. Datta, miR-539-5p regulates Srebf1 transcription in the skeletal muscle of diabetic mice by targeting DNA methyltransferase 3b. Mol Ther Nucleic Acids 29, 718–732 (2022).

63. F. Caiment et al., Assessing the effect of the CLPG mutation on the microRNA catalog of skeletal muscle using high-throughput sequencing. Genome Res 20, 1651–1662 (2010).

64. H. Takeda et al., The callipyge mutation enhances bidirectional long-range DLK1-GTL2 intergenic transcription in cis. Proc Natl Acad Sci U S A 103, 8119–8124 (2006).

65. S. K. Murphy et al., Callipyge mutation affects gene expression in cis: a potential role for chromatin structure. Genome Res 16, 340–346 (2006).

66. Y. Alkhaled et al., Impact of cigarette-smoking on sperm DNA methylation and its effect on sperm parameters. Andrologia 10.1111/and.12950 (2018).

67. H. Amor, Y. Alkhaled, R. Bibi, M. E. Hammadeh, P. M. Jankowski, The Impact of Heavy Smoking on Male Infertility and Its Correlation with the Expression Levels of the PTPRN2 and PGAM5 Genes. Genes (Basel) 14 (2023).

68. W. Zhang et al., CADM1 regulates the G1/S transition and represses tumorigenicity through the Rb-E2F pathway in hepatocellular carcinoma. Hepatobiliary Pancreat Dis Int 15, 289–296 (2016).

69. Y. Kasai et al., Trans-homophilic interaction of CADM1 promotes organ infiltration of T-cell lymphoma by adhesion to vascular endothelium. Cancer Sci 113, 1669–1678 (2022).

70. J. Dou, S. V. Thangaraj, Y. Zhou, V. Padmanabhan, K. Bakulski, Developmental programming: Differing impact of prenatal testosterone and prenatal bisphenol-A-treatment on hepatic methylome in female sheep. Mol Cell Endocrinol 609, 112655 (2025).

71. M. P. Wilson et al., A pseudoautosomal glycosylation disorder prompts the revision of dolichol biosynthesis. Cell 187, 3585–3601.e3522 (2024).

72. Y. Jiang et al., The sheep genome illuminates biology of the rumen and lipid metabolism. Science 344, 1168–1173 (2014).

73. M. Shihabi et al., Identification of Selection Signals on the X-Chromosome in East Adriatic Sheep: A New Complementary Approach. Frontiers in Genetics Volume 13 - 2022 (2022).

74. D. D. Jima et al., Genomic map of candidate human imprint control regions: the imprintome. Epigenetics 17, 1920–1943 (2022).

75. C. U. Braz et al., Paternal diet induces transgenerational epigenetic inheritance of DNA methylation signatures and phenotypes in sheep model. PNAS Nexus 1, pgac040 (2022).

76. J. Townsend, C. U. Braz, T. Taylor, H. Khatib, Effects of paternal methionine supplementation on sperm DNA methylation and embryo transcriptome in sheep. Environ Epigenet 9, dvac029 (2023).

77. M. Kizilaslan et al., Transgenerational Epigenetic and Phenotypic Inheritance Across Five Generations in Sheep. Int J Mol Sci 26 (2025).

78. K. D. Sinclair et al., Fetal growth and development following temporary exposure of day 3 ovine embryos to an advanced uterine environment. Reprod Fertil Dev 10, 263–269 (1998).

79. K. Gillespie, J. Stewart, Validation of iSperm analyzer for assessing ram semen quality. Proc Am Assoc Bovine Pract 58, (2024).

80. S. Parthipan et al., Spermatozoa input concentrations and RNA isolation methods on RNA yield and quality in bull (Bos taurus). Anal Biochem 482, 32–39 (2015).

81. A. Kozomara, M. Birgaoanu, S. Griffiths-Jones, miRBase: from microRNA sequences to function. Nucleic Acids Res 47, D155–d162 (2019).

82. Y. Chen, L. Chen, A. T. L. Lun, P. L. Baldoni, G. K. Smyth, edgeR v4: powerful differential analysis of sequencing data with expanded functionality and improved support for small counts and larger datasets. Nucleic Acids Res 53 (2025).

83. Y. Benjamini, Y. Hochberg, Controlling the False Discovery Rate: A Practical and Powerful Approach to Multiple Testing. Journal of the Royal Statistical Society: Series B (Methodological) 57, 289–300 (1995).

84. G. W. Montgomery, J. A. Sise, Extraction of DNA from sheep white blood cells. New Zealand Journal of Agricultural Research 33, 437–441 (1990).

85. H. Feng, H. Wu, Differential methylation analysis for bisulfite sequencing using DSS. Quant Biol 7, 327–334 (2019).

86. G. Yu, L. G. Wang, Q. Y. He, ChIPseeker: an R/Bioconductor package for ChIP peak annotation, comparison and visualization. Bioinformatics 31, 2382–2383 (2015).

87. Q. Wang et al., Exploring Epigenomic Datasets by ChIPseeker. Curr Protoc 2, e585 (2022)

