## Supplementary material for "Sex-specific multigenerational epigenetic responses to real-world chemical mixture exposure in an outbred sheep model": SI Appendix

**This PDF file includes:**

Supplementary Materials and Methods
Figures S1 to S9
Tables S1 to S6
SI References

**Other supplementary materials for this manuscript include the following:**

Datasets: File S1

#### Supplementary Information Text

**Materials and Methods**

All animal procedures were approved by the Animal Welfare and Ethical Review Board of the University of Glasgow. In addition, animal procedures were performed under the United Kingdom's Animals (Scientific Procedures) Act 1986. Associated protocols complied with the ARRIVE guidelines and with project licensed authority (PF10145DF).

**Experimental treatments, animals and design**

Multiparous EasyCare ewes (*Ovis aries*; F0, n = 320), that had not been exposed to BS previously, were allocated randomly to either Con or BS treated pastures at the University of Glasgow Cochno Farm and Research Centre. Specific BS treated pastures (2.25 tonnes dry matter BS per ha applied twice annually) have existed at this farm since 2014. Control pastures were treated with inorganic fertilizer (providing 225 kg nitrogen/ha/year), such that Con and BS treated pastures received equivalent amounts of nitrogen (Bellingham et al., 2009). Ewes were grazed on their respective pastures six weeks prior to mating (by artificial insemination; AI) until lambing, when all animals were turned out onto Con pastures and managed as a single flock thereafter.

Laparoscopic AI (Sinclair et al., 1998) was undertaken in 307 F0 ewes using fresh ejaculated semen collected from four unrelated (i.e., no common grandparents) F0 EasyCare rams (nominally identified in this article as A, B, C and D), that were not exposed to BS treated pastures, to generate four F1 (predominantly) half-sib family groups matched across the two treatment populations (Fig. S1 and Table S1) as reported previously (Evans et al., 2023). This served to standardize genetic variation between the two treatment groups within generation whilst avoiding the mating of close relatives. Subsequently, fresh semen collected from eight sexually mature (~18 months old) F1 rams (4 Con and 4 Biosolids), representing each of the four half-sib families within the two treatment groups, was used for laparoscopic AI of 149 age-matched sexually mature F1 ewes from the other three half-sib family groups. This ensured that the mating of close relatives was avoided and that the genetic structure of F2 offspring were closely matched for Con and BS treatment groups (Fig. S1 and Table S2). Finally, natural matings between available sexually mature, age-matched F2 rams and ewes (n = 61) were undertaken representing and matching the full range of available families within each of the two treatment groups (Fig. S1 and Table S3).

For semen analyses in the current article, additional collections were taken from a subset of 16 F1 and 14 F2 sexually mature rams on two occasions within generation, ensuring that F1 and F2 genetic backgrounds were balanced between treatment groups (Table S1). Ejaculated semen was processed immediately for analysis and cryopreservation.

**Computer assisted sperm assessment (CASA)**

A 50 µL aliquot of fresh semen was used for concentration and motility analysis using the iSperm mCASA semen analyzer (Aidmics Biotechnology Co., Ltd, Taipei, Taiwan). Optimized across species including sheep, some reports indicate that concentration measurements are more variable than motility measures (Dini et al., 2019, Bulkeley et al., 2021, Gillespie and Stewart, 2024). Consequently, when diluting to between 30 and 60M/mL (the iSperm working range), a 1:100 dilution was made in water to immobilize for counting, and in INRA 96 semen extender (Elite Reproduction Services, Shifnal, UK) for motility analysis. In both analyses, a 7.5ul volume of thoroughly mixed diluted ejaculate was pipetted onto the surface of the base chip mounted onto the iSperm light source. The base chip was inverted, placed into a cover chip vertically and attached to a mini-iPad camera with temperature maintained at 37˚C. Sperm concentration and motility were recoded Using the iSperm Ovine 5 app (Gillespie and Stewart, 2024). Parameters analyzed included motility, progressive motility and straight-line velocity, and were based on triplicate measurements per sample. The remaining ejaculate was transferred to a 1.5 mL Eppendorf vial and centrifuged at 16,000g and 4˚C for 20 min, with the supernatant centrifuged for a second time. Both the supernatant (seminal plasma) and residual sperm were snap frozen in liquid nitrogen and stored at -80˚C until analysis.

**Sperm preparation for RNA extraction**

Frozen sperm pellets were suspended in cold PBS and 500 million sperm purified in 60% BoviPure (Nidocon, Sweden) diluted in PBS. Following centrifugation (300g for 25 min at 4°C), washed pellets were re-suspending in 4 ml lysis buffer (0.1% SDS and 0.5% Triton x-100 in molecular biology grade water). Sperm suspensions were incubated on ice for 30 min before centrifugation at 800 g for 5 min at 4°C. Pellets were washed with PBS again and a suspension equivalent to 100 million sperm aliquoted into 2 mL LoBind tubes and spin at 12000 g for 5 min at 4°C before snap freezing in liquid nitrogen and storage at -80°C.

**RNA extraction from sperm**

RNA was extracted using miRNeasy kit (Qiagen) (Parthipan et al., 2015) with a slight modification. Briefly, 1.6 mL of RLT (Qiagen) containing beta-mercaptoethanol was added. Pellets were homogenized by passing through a 19 g needle fitted to 5 ml syringe 8 times and incubated at room temperature for 20 min. Each lysate was split into 8 x 200 µL and 1 mL QIAzol (Qiagen) was added per 200 µL lysate, mixed and incubated at room temperature for 5 min. Then 200 µL chloroform was added to each tube and mixed vigorously by hand for 30 sec and tubes left at room temperature for 5 min. Mixtures were spun at 12,000xg for 15 min at 4°C for phase separation. After centrifugation, the top phase was transferred and 1.5 volume of 100% ethanol added. Mixtures were mixed by inverting tubes. RNA was extracted using RNeasy miRNA kit (Qiagen) according to manufacturer's instructions and on column DNase treatment was included in the extraction. Samples were quantified by Qubit miRNA kit.

**RNA extraction from seminal plasma**

RNA was extracted from 200 µL seminal plasma using miRNeasy mini kit (Qiagen, UK) as described by manufacturer. Briefly, frozen seminal plasma was thawed at room temperature. Then 1 ml of QIAzol (Qiagen, UK) was added to 200 µl of thawed seminal plasma, mixed and incubated at room temperature for 5 min. After addition of 200 µl of chloroform, the mixture was mixed vigorously for 30 sec and left at room temperature for 3 min, followed by centrifugation at 12,000 g for 15 min at 4°C. The upper phase was mixed with 1.5 volume of absolute ethanol. This mixture was transferred to RNeasy MinElute spin columns (RNeasy MinElute Cleanup kit, Qiagen, UK). Following washing with buffers RWT and RPE from the kit, columns were washed once with 80% ethanol before eluting in 15 µl of RNase-free water. miRNA concentration was quantified using Qubit miRNA kit (Invitrogen Ltd, Paisley, UK).

**miRNA library preparation and sequencing**

Indexed miRNA libraries were prepared using the QIAseq miRNA Library Prep kit (Qiagen; 331505) and the QIAseq miRNA 96 Index Kit IL-UDI-A (Qiagen; 331615). The amount of input RNA per sample was 25 ng and 17 cycles of PCR were used for library amplification. Libraries were quantified using the Qubit Fluorometer and the Qubit dsDNA HS Kit (ThermoFisher Scientific; Q32854). Library fragment-length distributions were analyzed using the Agilent 4200 TapeStation and the Agilent High Sensitivity D1000 ScreenTape Assay (Agilent; 5067-5584 and 5067-5585). Libraries were pooled in equimolar amounts and final-library QC was performed using the Qubit Fluorometer and the Qubit dsDNA HS Kit and the TapeStation High Sensitivity D1000 ScreenTape Assay. The library pool was sequenced on the Element Biosciences AVITI System on an AVITI 2X75 Sequencing Kit - Cloudbreak FS Medium Output (Element Biosciences; 860-00014), generating approximately 20 million 75-bp single-end reads per sample.

**DNA extraction**

Genomic DNA was extracted from liver tissue using the DNeasy Blood and Tissue Kit (Qiagen), following the manufacturer's protocol. RNase A treatment was included to eliminate residual RNA contamination. Sperm that had been through gradient and somatic cell lysis was treated with DNase to remove cell-free DNA before DNA extraction. Pellets containing 10 million sperm were resuspended in 87.5 µL PBS, 10 µL RDD buffer (Qiagen, Manchester, UK) and 2.5 µL DNase I solution (Qiagen). Samples were incubated at room temperature for 10 min and centrifuged at 16,000xg for 10 min at 4°C. Pellets were then washed twice with PBS before subjected to DNA extraction using DNeasy kit (Qiagen) with modification from user developed protocol 1 QA03 (Qiagen). To DNase-treated pellets, 250 µL lysis buffer (100 mM Tris-HCl, 10 mM EDTA, 500 mM NaCl, 1% SDS, 2% β-mercaptoethanol) and 50 µL proteinase K (Qiagen) were added and incubated at 56°C for 2 h. After 2 h another 20 µL of proteinase K was added and samples were incubated for a further 2 h at 56°C. Samples were then purified using DNeasy kit with on column RNase treatment during extraction. DNA was eluted in EB buffer and quantified using Nanodrop. DNA integrity was checked on 1.2% agarose gel. Red blood cells were lysed from peripheral blood and the remaining white blood cells pelleted by centrifugation at 2,500g for 10 min at 10°C. Genomic DNA was extracted using the salting-out method (Montgomery and Sise, 1990), with minor modifications. Briefly, cells were lysed and digested overnight at 55°C in 800 µL of lysis buffer containing 10 mM Tris-HCl (pH 8.0), 0.1 mM EDTA, 0.5% sodium dodecyl sulfate (SDS), and 0.1 mg/mL proteinase K. The following morning, an additional 30 µg of proteinase K was added, and samples were incubated for a further 3 hours at 55°C to ensure complete protein digestion. Subsequently, 360 µL of 5 M sodium chloride (NaCl) was added to each sample, followed by vigorous shaking for 15 s. Samples were incubated at room temperature for 10 min and centrifuged at 12000g for 5 min to pellet the precipitated proteins. The supernatant was transferred to a new tube, and genomic DNA was precipitated by adding two volumes of absolute ethanol. Tubes were inverted several times until visible DNA strands formed. The DNA strands were transferred to a new tube and washed twice with 70% ethanol, followed by centrifugation at 12000g for 2 min. The resulting DNA pellet was air-dried and resuspended in 100 µL of TE buffer (10 mM Tris-HCl, 1 mM EDTA, pH 8.0). Samples were incubated at 4°C overnight to ensure DNA was completely dissolved. DNA concentration and purity were assessed using a DeNovix spectrophotometer (DeNovix, USA).

**Reduced Representation Bisulfite Sequencing**

DNA from sperm, blood and liver was diluted to 15 ng/µl with water and concentration was checked with Qubit dsDNA HS kit (ThermoFisher, Loughborough, UK). Then 100 ng of DNA and 800 pg unmethylated λ (Promega, Southampton, UK) were treated with 20 units *Msp1* enzyme (NEB, Hitchin, uk), followed by end repair and A-tailing. Methylated adaptor from NEBNext Multiplex Oligos for enzymatic methyl-seq kit (NEB, E7140L) was then ligated to DNA using Blunt end/TA ligase master mix (NEB) according to manufacturer's instruction. Adaptor-ligated DNA samples were cleaned up using Zymo clean and concentrator before bisulfite conversion using EZ methylation gold kit (Zymo, CA, USA). Following completion of bisulfite conversion cycle and purification, RRBS libraries were prepared using bisulfite-converted DNA, KAPAHiFi Uracil + mix (Roche Diagnostics Ltd, West Sussex, UK) and dual index primer pairs (NEB, E7140L). PCR reaction was performed in 50 uL at 98°C for 45 sec followed by 8 cycles of 98°C for 15 sec, 60°C for 30 sec and 72°C for 30 sec and single cycle of 72°C for 1 min. PCR products were then purified using MinElute PCR kit (Qiagen). Equimolar amounts of libraries were pooled and 200-600 bp were size-selected using BluePippin (Sage Science, Beverly, MA, USA). Size-selected fragments were checked by Qubit and Bioanalyser. Pooled libraries were sequenced in a single lane with 2% Phix spike in using Illumina NovoSeq X plus to give at least 80 million 150 bp pair-end reads per sample (Novogene, Cambridge, UK). This equated to an average read depth of ≥ 25 reads per cytosine for blood and liver DNA across libraries and ≥ 35 reads per cytosine for sperm DNA libraries.

**miRNA Read Handling and Differential Expression**

Raw sequencing reads were run through the Nextflow nf-core/smrnaseq pipeline v2.3.1 (Di Tommaso et al. 2017; Ewels et al. 2020) and aligned to 153 mature sequences arising from 106 precursor sheep miRNA sequences from the miRBase database (Kozomara et al. 2019) using Bowtie1 (Langmead et al. 2009). Differential expression (DE) analyses were then performed using the Bioconductor edgeR package v4.0.16 (Chen et al. 2025), looking for differences in BS exposed rams compared to Con, accounting for sire as a batch effect in analyses of all samples. miRNAs filtered to retain miRNAs with at least 10 counts per million in ≥ 50% of samples. Normalized counts were fitted in a negative binomial generalized linear model (nbGLM) using the genewise quasi-likelihood F-tests method. Exploratory F2 single sire analyses instead used a conservative, estimated dispersion value, with counts then fitted to a nbGLM using genewise likelihood ratio tests. DE miRNAs were defined as those with a log fold change ≥ 2.0 and an observed p-value ≤ 0.05. The Benjamini–Hochberg procedure (Benjamini and Hochberg, 1995) accounted for multiple testing, with a FDR ≤ 0.01 considered significant.

**Differential Methylation Analyses**

Sequenced RRBS libraries were initially run through the Nextflow nf-core/methylseq pipeline v3.0.0 (Di Tommaso et al. 2017; Ewels et al. 2020; Ewels et al. 2024) with default options apart from the RRBS option being enabled, trimming options set to trim the first 3 bases of all reads and the aligner set to bismark for aligning processed reads to the Ensembl reference genome ARS-UI_Ramb_v2.0 (GCA_016772045.1). Generated methylation coverage files were filtered to remove methylated CpGs (mCpGs) matching positions of known SNPs and then masked using methrix (Mayakonda et al. 2020) to remove mCpGs with coverage less than five, or in the highest 1% of coverage, and to include only mCpGs present within 75% of samples per treatment group. Differential methylation analyses were then performed using DSS (Feng & Wu 2019). DMLs were identified using the DMLtest() function with default options and smoothing enabled, and the callDML() function with a delta of 0.15 (≥ 15% difference in methylation) and a p threshold of 0.05. The Benjamini–Hochberg procedure (Benjamini and Hochberg, 1995) accounted for multiple testing, with a FDR ≤ 0.01 considered significant.

Once identified, DMLs were labelled genic or intergenic using the annotatePeak() function from the ChIPseeker v1.34.1 package (Yu et al. 2015; Wang et al. 2022). The Bioconductor topGO v2.54.0 package (Alexa and Rahnenfuhrer 2023) was then used to perform gene ontology (GO) analyses on genic DML using Fisher's Exact Test with the Elim algorithm. A node size of 5 was used to prune GO terms with less than the specified number of annotated genes. KEGG enrichment analyses were performed using the enrichKEGG() function from clusterProfiler v4.6.2 (Yu et al. 2012; Wu et al. 2021) with a p-value cutoff of 0.05. Additionally, genes of interest that may escape genome-wide demethylation in embryonic development were identified from a list of pig and human 'escapee' orthologs (Zhu et al. 2021), and their enrichment determined using Fisher's exact test.

**Quantitayive Real-Time PCR (qPCR)**

Total RNA was isolated from liver using the RNeasy Mini Kit (Qiagen) with on-column DNase I treatment to remove genomic DNA, as per the manufacturer's instructions. RNA was eluted in 50 µL of RNase-free water, and its concentration and purity were assessed using a DeNovix spectrophotometer. Reverse transcription was performed using 900 ng of total RNA with the QuantiTect Reverse Transcription Kit (Qiagen), according to the manufacturer's protocol. The resulting cDNA was diluted 1:15 with RNase-free water prior to use in qPCR, which was conducted using the QuantiNova SYBR Green PCR Master Mix (Qiagen) on a Bio-Rad CFX Opus Real-Time PCR System (Bio-Rad Laboratories Ltd, Hemel Hempstead, UK). Reactions were undertaken in 384-well plates with a final volume of 12 µL, comprising 6 µL of SYBR Green mix, 2 µL of diluted cDNA, and 0.4 µL each of 10 µM forward and reverse primers (see table below). RNase-free water was used as a negative control. Thermal cycling conditions were as follows: initial enzyme activation at 95°C for 2 min, followed by 40 cycles of denaturation at 95°C for 5 sec and annealing/extension at 60°C for 10 sec with data acquisition. A melt curve analysis was performed from 60°C to 95°C in 0.5°C increments with a 5-sec hold at each step. Primers (Table Si) were designed using Primer-BLAST (NCBI, USA) to span exon-exon junctions, ensuring specificity for cDNA. Primer specificity was validated by sequencing the PCR products.

#### Statistical analysis

Analyses were performed using the GenStat statistical package (21st Edition, VSN International, 2022; https://www.vsni.co.uk/). Sperm motility (CASA) and relative transcript expression (F3 livers) were analyzed using restricted maximum likelihood (REML) generalized linear mixed models. The fixed term fitted to these models was treatment, with sire group (F1) or grandsire group (F2) fitted as random terms for sperm analysis; and great grandsire fitted as the random term for F3 transcript analysis. Data are presented as predicted means ± SEM.

**Table Si.** Primer design for qPCR.

| Gene | Accession no. | Primers sequence 5' to 3' | Product length |
|---|---|---|---|
| Cell adhesion molecule 1 (CADM1) | XM_060399309.1 | F: GGCCCAGCCTGTTCATCAAT<br>R: TCACCTGCTCGAGAATCGTAT | 127 |
| H2A.Z variant histone 1 (H2AZ1) | NM_001009270.1 | F: AGTGGGCCGTATTCATCGAC<br>R: AGTGGGCCGTATTCATCGAC | 123 |
| Hypoxanthine phosphor-ribosyl-transferase 1 (HPRT) | XM_015105023.3 | F: ACTGAAGAGCTACTGTAACGACC<br>R: GACCAAGGAAAGCAAAGTCTG | 148 |
| Peptidylprolyl isomerase A (PPIA) | NM_001308578.1 | F: TCCCGAAGACAGCAGAAAACT<br>R: GGACTTGCCACCAGTACCAT | 146 |
| Succinate dehydrogenase complex flavoprotein subunit A (SDHA) | XM_027980212 | F: ACTACAAGGGGCAGGTTCTG<br>R: GAGTTTGCACCCAGACGGTT | 121 |

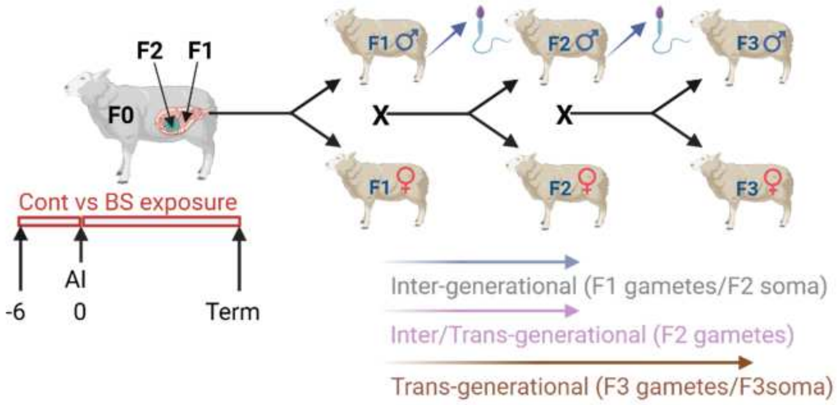


| Sire | A | B | C | D | A | B | C | D |
|---|---|---|---|---|---|---|---|---|
| Treatment | Control (Con) | | | | Biosolids (BS) | | | |
| F1 | A | B | C | D | A | B | C | D |
| F2 | AB AC AD BA BC BD | | | | AB AC AD BA BC BD | | | |
| | CA CB CD DA DB DC | | | | CA CB CD DA DB DC | | | |
| F3 | ABCD | BACD | CABD | DABC | ABCD | BACD | CABD | DABC |
| | ABDC | BADC | CADB | DACB | ABDC | BADC | CADB | DACB |
| | ACBD | BCAD | CBAD | DBAC | ACBD | BCAD | CBAD | DBAC |
| | ACDB | BCDA | CBDA | DBCA | ACDB | BCDA | CBDA | DBCA |
| | ADBC | BDAC | CDAB | DCAB | ADBC | BDAC | CDAB | DCAB |
| | ADCB | BDCA | CDBA | DCBA | ADCB | BDCA | CDBA | DCBA |

**Fig. S1.** Experimental design and breeding schedule. F0 ewes were exposed to Control (Con) or Biosolids (BS) treated pastures from 6 weeks prior to pregnancy establishment (by laparoscopic artificial insemination (AI)) and throughout gestation. BS exposure was terminated at parturition. As depicted, gestating F0 ewes carried F1 fetuses to term. These fetuses harbored germ cells that give rise to F2 individuals. For initial pregnancy establishment, semen was collected from 4 unrelated (with no-common parents or grandparents) F0 sires (nominally identified as A, B, C and D), that were not exposed to BS treated pastures. Unrelated F0 ewes of the same breed (EasyCare), that had not previously been exposed to BS treated pastures, were allocated at random to each treatment and to each sire. This generated 4 half-sib F1 family groups matched across treatments. Sexually mature F1 ewes were also bred by laparoscopic AI to generate F2 offspring using semen from 4 Con and 4 BS exposed sexually mature rams from each of the 4 half-sib families to impregnate F1 ewes from each of the other three half-sib family groups within treatment. Adopting the same philosophy of breeding between families within treatment groups, available male and female F2 offspring were, on this occasion, naturally mated to generate F3 offspring. This breeding schedule ensured that the mating of close relatives was avoided and that genetic structure within generation was matched between the two BS treatment groups. Liver samples were collected from each of the three generations and, in addition, whole blood from F2 offspring, all for DNA methylation analyses. Semen from F1 and F2 males was also collected for epigenetic analyses (i.e., microRNA and DNA methylation).

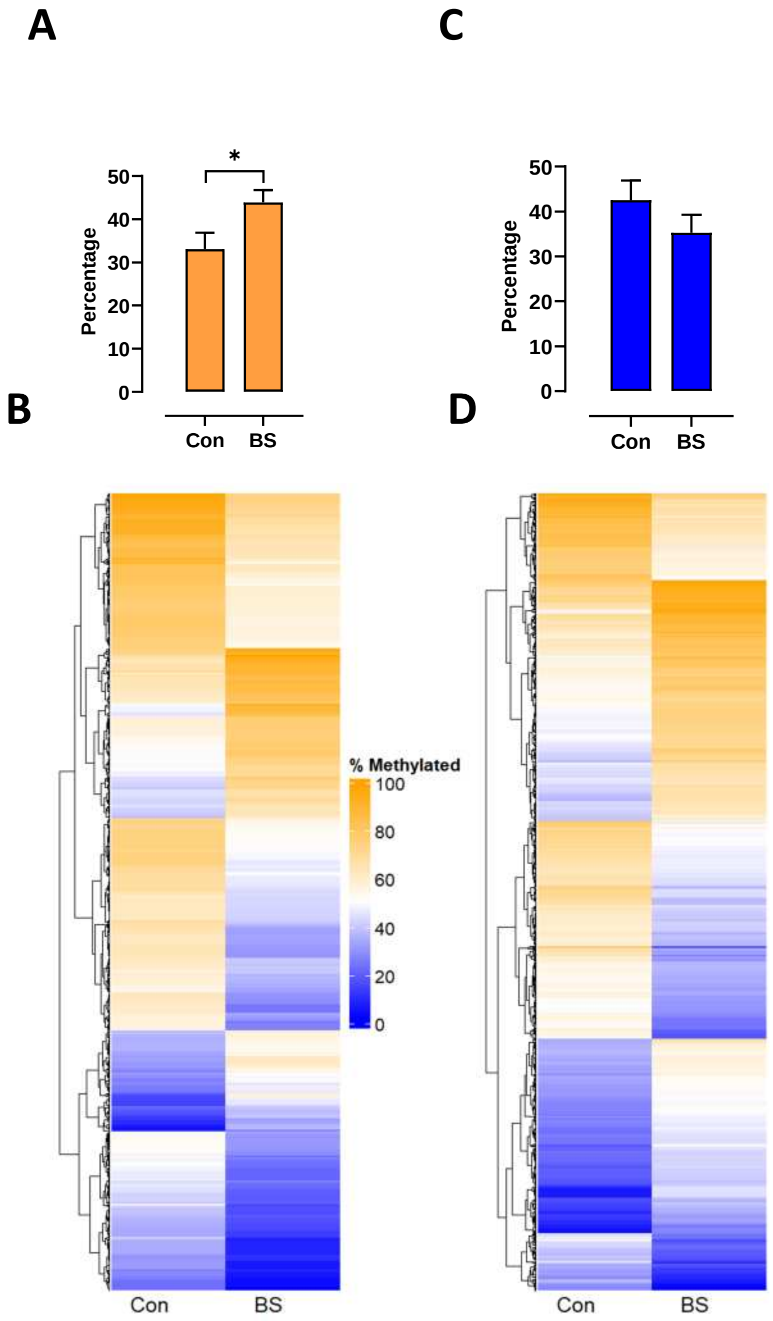


**Fig. S2.** Physiological and molecular characteristics of F1 and F2 semen (i.e., sperm and seminal plasma). **A.** Mean ± SEM percentage progressive sperm motility in F1 rams (from Sires A, B, C and D) established using the iSperm for Control (Con) and Biosolids (BS) experimental groups; **B.** Heatmap highlighting mean (across the four sire families) differentially methylated loci (DML; 15% threshold) in F1 sperm for Con and BS experimental groups; **C.** Mean ± SEM percentage progressive motility in F2 rams established using the iSperm for Con and BS experimental groups; **D.** Heatmap highlighting mean (across all grandsire families) DML in F2 sperm for Con and BS experimental groups.

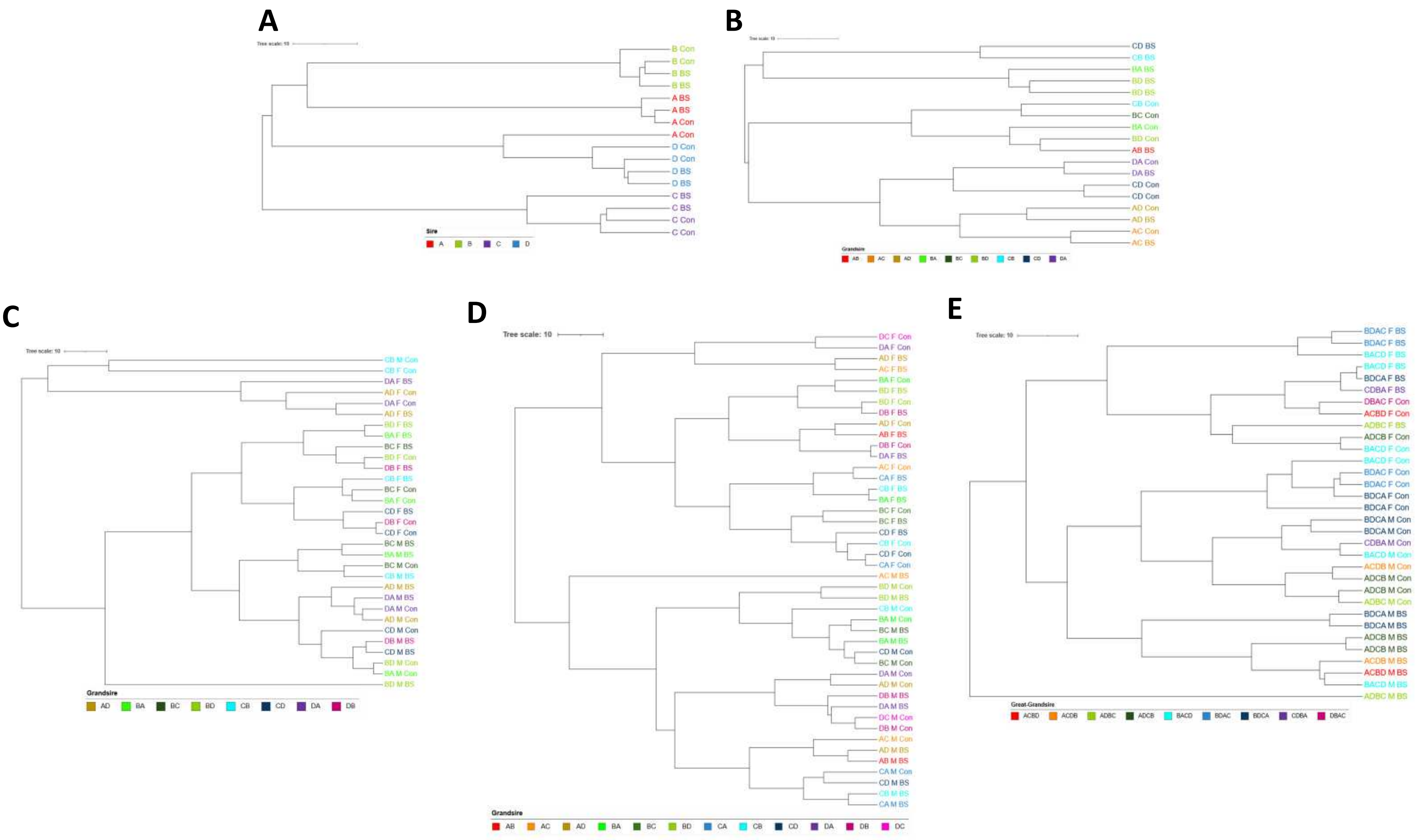


**Fig. S3.** Hierarchical clustering of DNA methylation. Distances calculated from top 3 principal components. Label colours indicate sire/grandsire/great-grandsire groupings. **A.** F1 sperm. **B.** F2 Sperm. **C.** F2 liver. **D.** F2 blood. **E.** F3 liver.

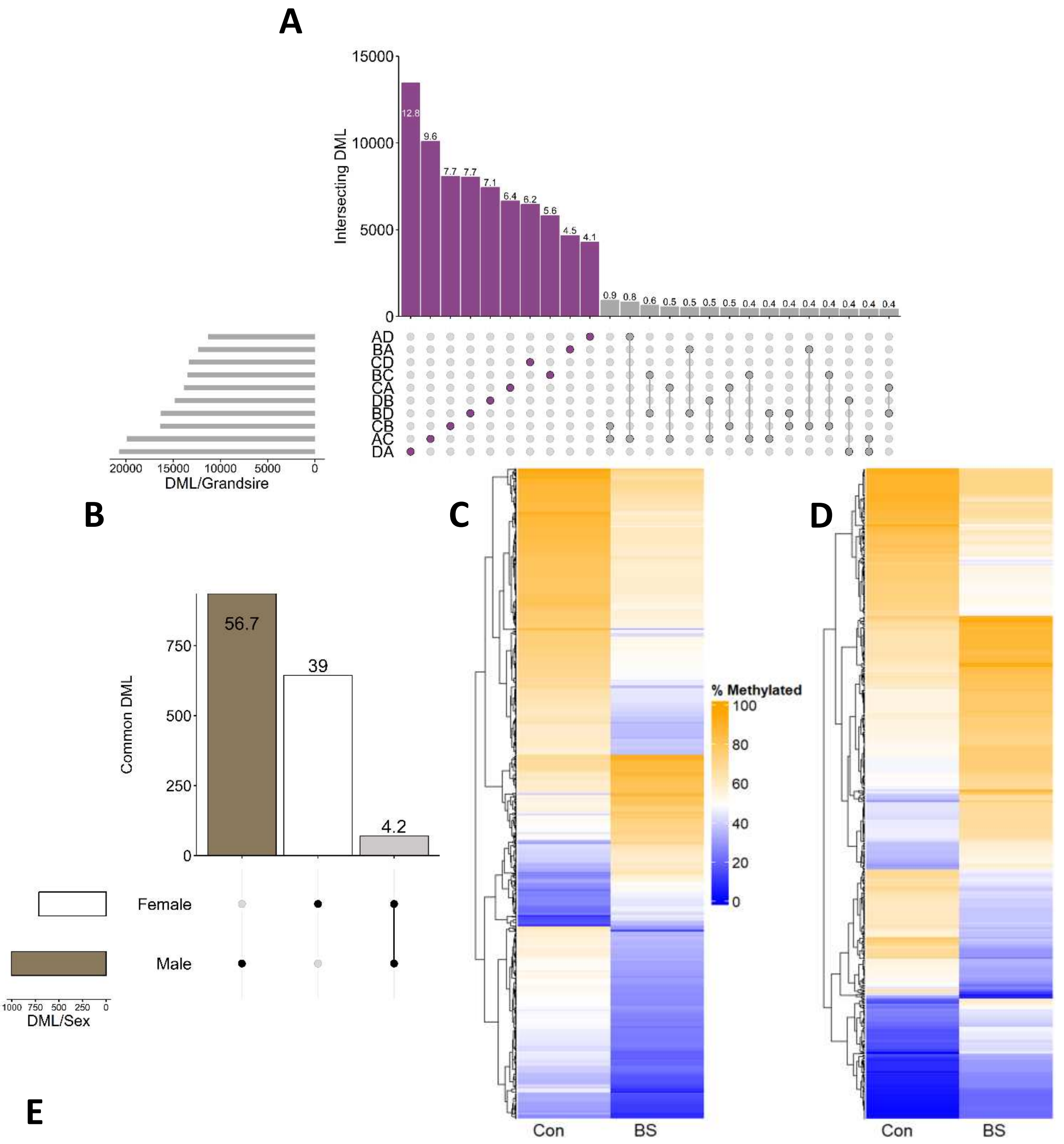


E

| | Sire AC | Sire AD | Sire BA | Sire BC | Sire BD | Sire CA | Sire CB | Sire CD | Sire DA | Sire DB | Male | Female |
|---|---|---|---|---|---|---|---|---|---|---|---|---|
| DML | 19,923 | 11,305 | 12,337 | 13,503 | 16,322 | 13,859 | 16,404 | 13,319 | 20,729 | 14,846 | 1,006 | 714 |
| Genes | 2,108 | 1,191 | 1,323 | 1,449 | 1,735 | 1,557 | 1,856 | 1,435 | 1,657 | 1,670 | 111 | 86 |

**Fig. S4.** Methylation Characteristics of F2 blood. **A.** Upset plot of overlapping differentially methylated loci (DML) (15% threshold) across the seven F2 grandsire groups (Blue: grandsire specific, Grey: overlapping; values on bars denote percentage of total DML). Only the top 25 intersections are displayed. **B.** Upset plot of overlapping DML across male and female F2 liver (Black/White: sex specific, Grey: overlapping; values on bars denote percentage of total DML). **C.** Heatmap highlighting mean methylation of DML in males in the Control (Con) and Biosolids (BS) exposed experimental groups. **D.** Heatmap highlighting mean methylation of DML in females in the Con and BS exposed experimental groups. **E.** DML and DML-containing genes counts across ten grandsire groups, as well as in males and females.

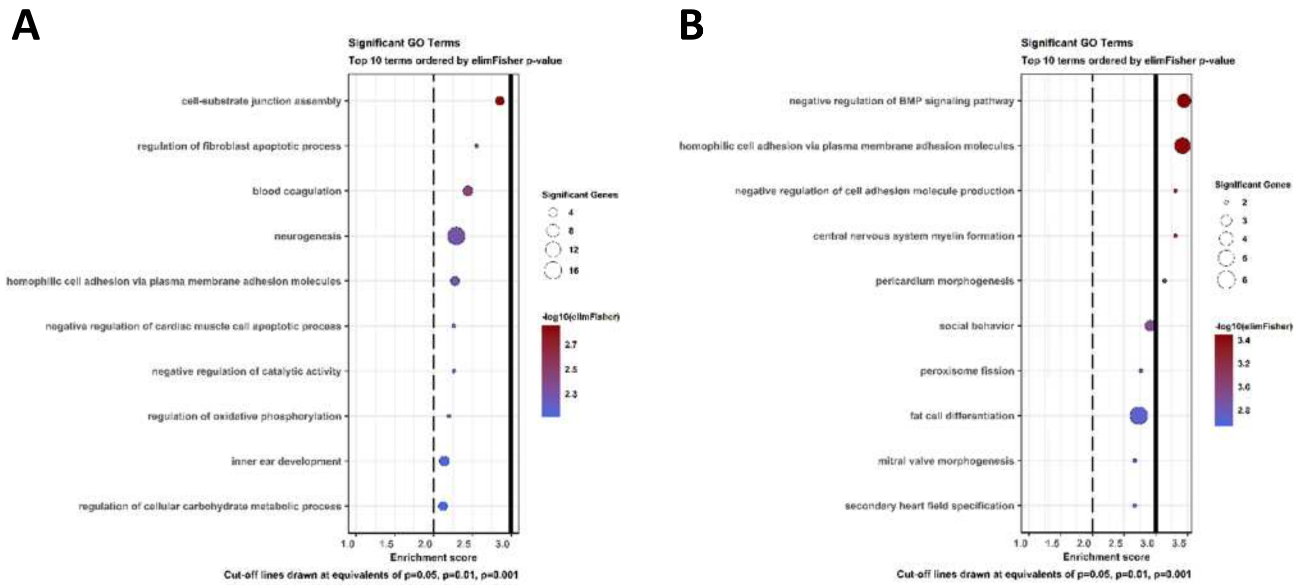


**Fig. S5.** Top 10 biological GO terms derived from genes containing DMLs within F3 livers. **A.** Males. **B.** Females.

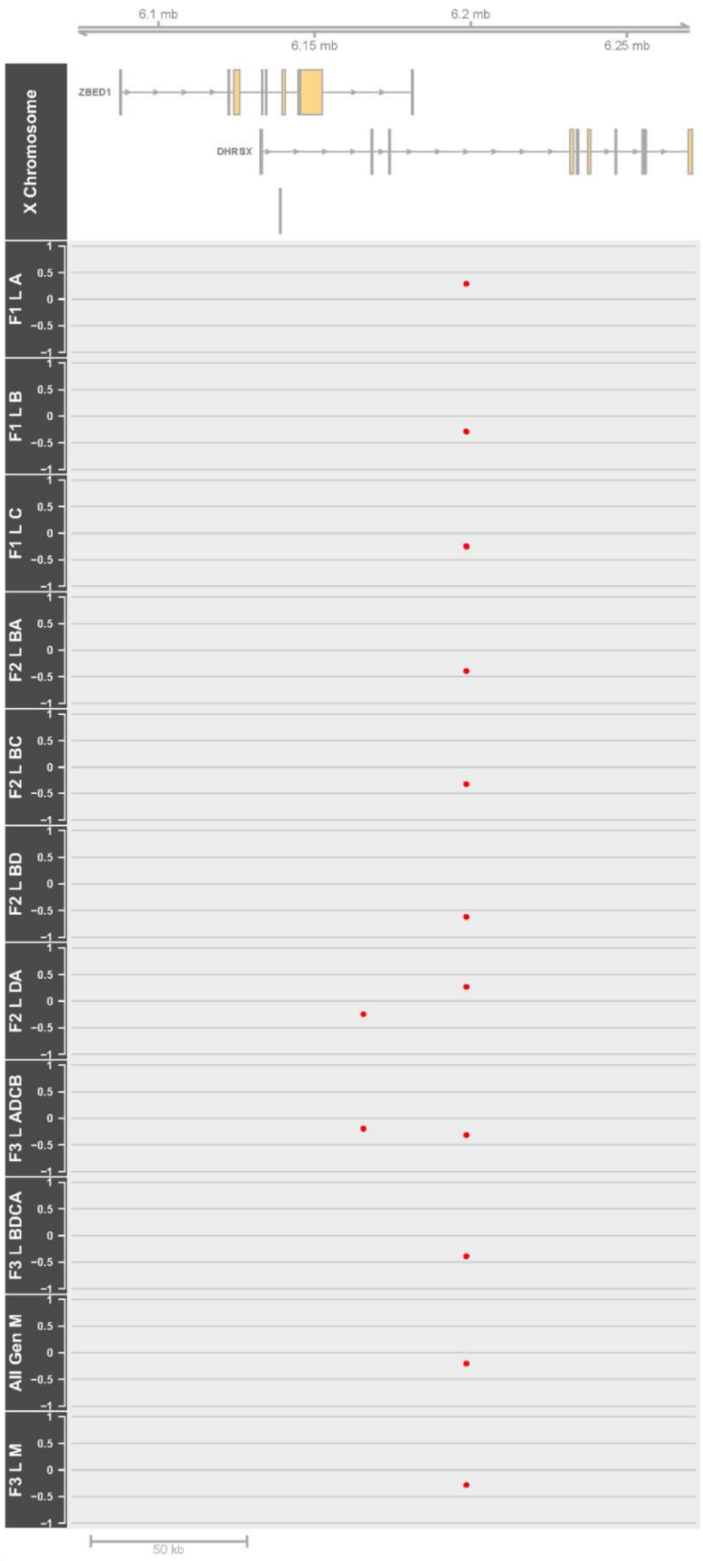


**Fig. S6A.** Genomic locations of cross-generational DML in sire/grandsire/great-grandsire groups in F1, F2 and F3 generations within *DHRSX.* Methylation differences in biosolids-exposed animals compared to control are displayed per group. 7 DML within *DHRSX* Intron 3, identified in each generation of lineages A, B and C, as well as in F3 males and males when all three generations of liver samples were analyzed together.

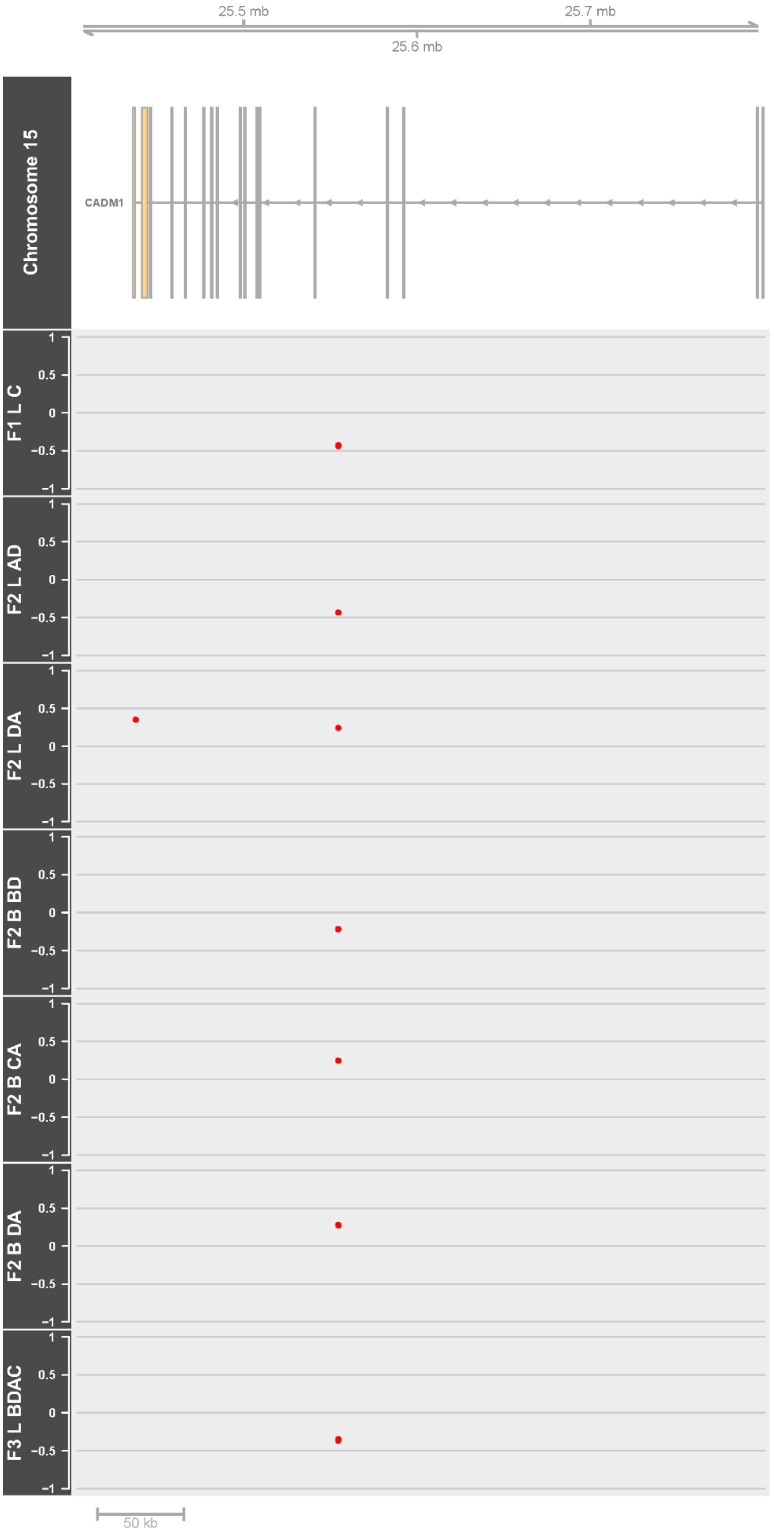


**Fig. S6B.** Genomic locations of cross-generational DML in sire/grandsire/great-grandsire groups in F1, F2 and F3 generations within *CADM1*. Methylation differences in biosolids-exposed animals compared to control are displayed per group. A. 10 DML identified within at least one sire/grandsire/great-grandsire group per generation within *CADM1*, Intron 4 in liver. F2 blood grandsires containing these DML are also shown. "L" = Liver; "B" = Blood.

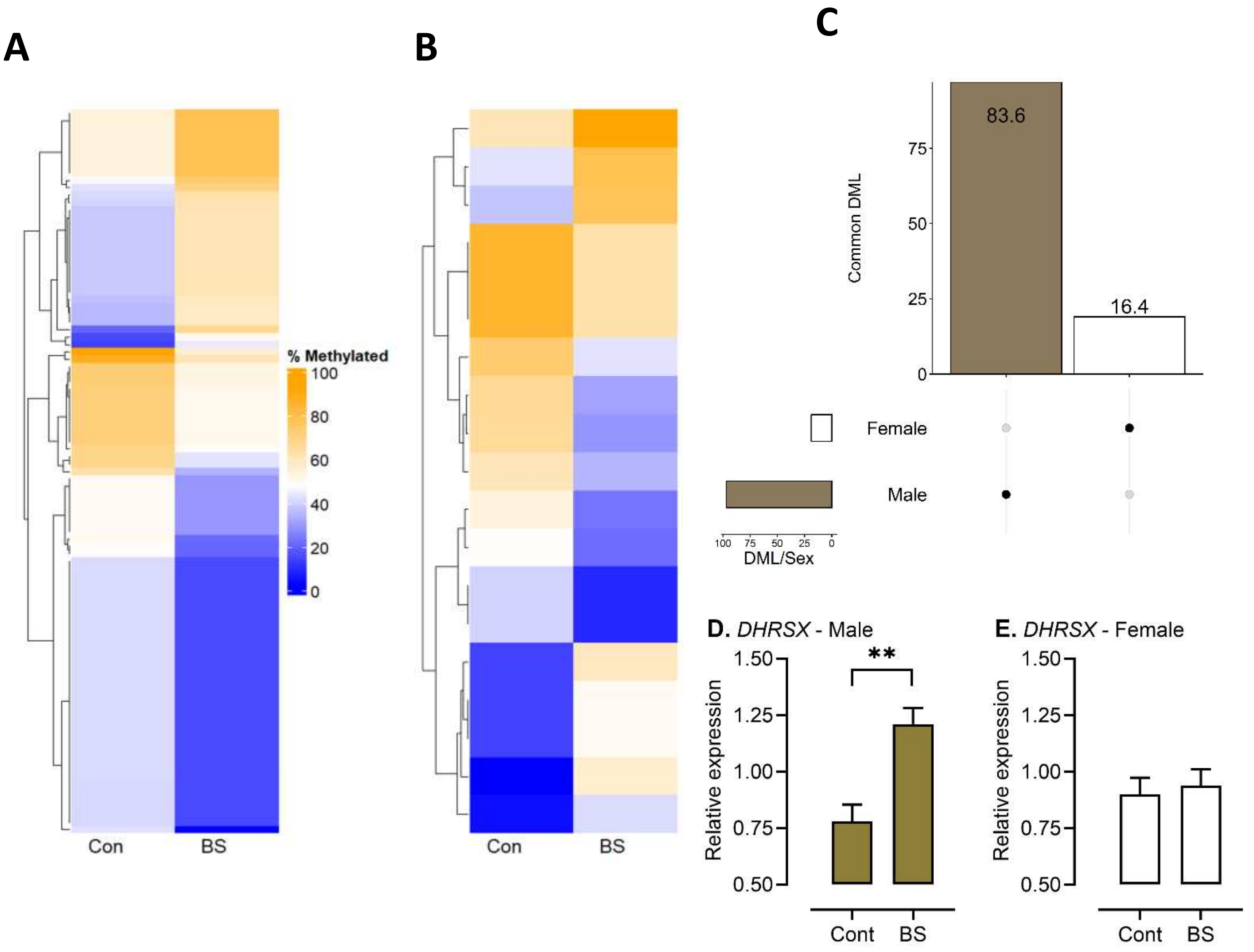


**Fig. S7.** Cross-generational differential methylation analyses run on liver samples from all generations, averaged across sire/grandsire/great-grandsire and generation, split by sex. **A.** Cross-generational differential methylated loci (DML) in males. **B.** Cross-generational DML in females. **C.** Intersecting cross-generational DMLs across males and females (Brown/White: sex specific; values on bars denote percentage of total DML). **D.** DHRSX expression in Control (Con) and Biosolids (BS) exposed male F3 liver (** $P < 0.01$). **E.** DHRSX expression profile in Con and BS exposed female F3 liver.

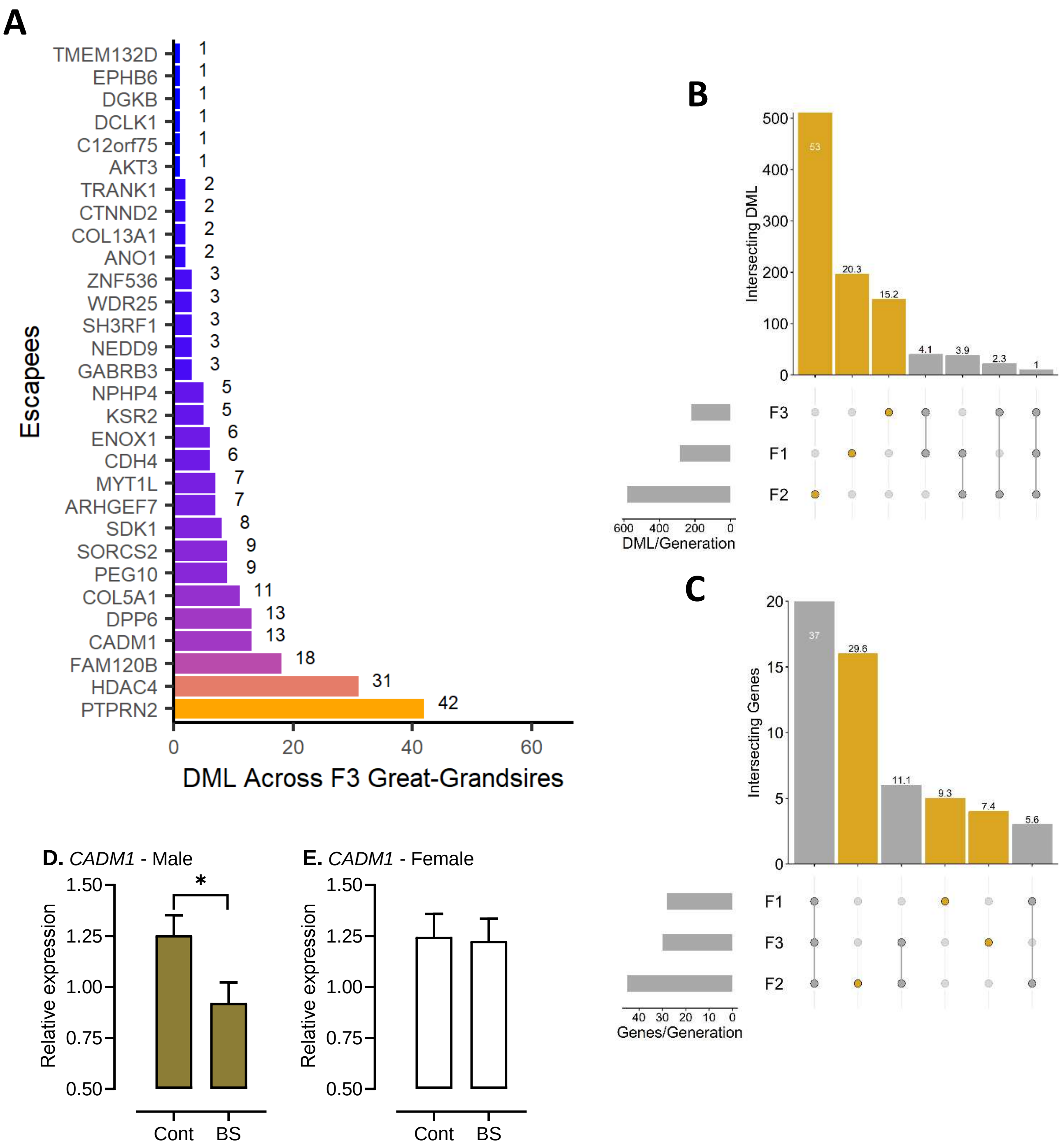


**Fig. S8.** F3 hepatic differentially methylated loci (DML) (15% threshold) in genes predicted (Zhu et al., 2021) to escape genome wide demethylation during embryonic development ('escapees'). **A.** Total unique genic escapee DML from single great-grandsire analyses. **B.** Intersecting genic DML (split by sire, grandsire, and great-grandsire, and averaged across sex) within 'escapees' in liver samples across 3 generations of offspring. **C.** Intersecting 'escapees' containing DML in liver samples across 3 generations of offspring. **D.** CADM1 expression in Control (Con) and Biosolids (BS) exposed male F3 liver. **E.** CADM1 expression profile in Con and BS exposed female F3 liver.

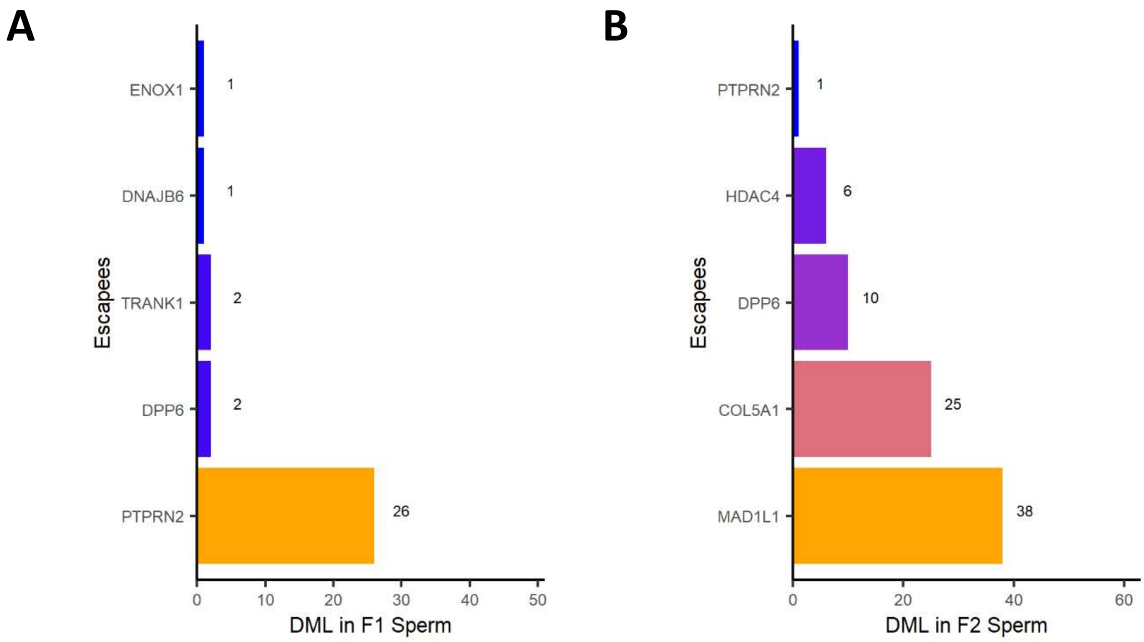


**Fig. S9.** Sperm differentially methylated loci (DML) (15% threshold) in genes predicted (Zhu et al., 2021) to escape genome wide demethylation during embryonic development ('escapees'). **A.** Genic 'escapee' DML from F1 sperm. **B.** Genic 'escapee' DML from F2 sperm.

**Table S1.** F0 Pregnant and lambed (F1) following laparoscopic AI.

| | **Con** | **BS** | **Significance** |
|---|---|---|---|
| Ewes inseminated, n | 156 | 151 | |
| Pregnant at scanning, % | 82.7 ± 3.00 | 79.3 ± 3.27 | - |
| Ewes lambed, % | 75.7 ± 3.40 | 69.9 ± 3.70 | - |
| Lambed of scanned pregnant, % | 89.9 ± 2.21 | 87.4 ± 2.52 | - |
| Litter size at lambing, n | 2.04 ± 0.167 | 1.96 ± 0.156 | - |
| Lambs of ewes inseminated, n | 1.43 ± 0.096 | 1.33 ± 0.094 | - |

**Table S2.** F1 pregnancy and lambed (F2) following laparoscopic AI.

| | Con | BS | Significance |
|---|---|---|---|
| Ewes inseminated, n | 77 | 72 | |
| Pregnant at scanning, % | 67.8 ± 5.04 | 70.5 ± 5.14 | - |
| Ewes lambed, % | 61.5 ± 5.21 | 64.8 ± 5.29 | - |
| Lambed of scanned pregnant, % | 91.6 ± 2.72 | 90.8 ± 3.06 | - |
| Litter size at lambing, n | 1.56 ± 0.187 | 1.63 ± 0.191 | - |
| Lambs of ewes inseminated, n | 0.92 ± 0.027 | 0.91± 0.031 | - |

**Table S3.** F2 pregnancy and lambed (F3) following natural mating.

| | Con | BS | Significance |
|---|---|---|---|
| Ewes inseminated, n | 27 | 34 | |
| Ewes lambed, % | 70.6 ± 9.02 | 73.2 ± 7.21 | - |
| Litter size at lambing, n | 1.74 ± 0.302 | 1.72 ± 0.262 | - |
| Lambs of ewes inseminated, n | 1.22 ± 0.213 | 1.27 ± 0.193 | - |

**Table S4.** Live born F1 progeny split by treatment, sex and sire-family depicting, in parentheses, samples collected for epigenetic analyses.

| **Treatment** | **Control** | | **Biosolids** | | **Total** |
|---|---|---|---|---|---|
| **Sex** | **M** | **F** | **M** | **F** | |
| **Sire family groups** | | | | | |
| A | 27 (1) (2) | 24 (1) | 29 (1) (2) | 22 (1) | **102** |
| B | 32 (1) (2) | 23 (1) | 25 (1) (2) | 22 (1) | **102** |
| C | 28 (1) (2) | 28 (1) | 26 (1) (2) | 28 (1) | **110** |
| D | 16 (1) (2) | 23 (1) | 25 (1) (2) | 23 (1) | **88** |
| **Total** | **104** | **98** | **105** | **95** | **402** |

Numbers in blue parentheses depict 16 liver samples from 8-week-old F1 offspring used for RRBS. Numbers in red parentheses depict 16 sperm and seminal plasma samples from sexually mature F1 offspring used for epigenetic analyses

**Table S5.** Live born F2 progeny split by treatment, sex and grandsire family

| Treatment | Control | | Biosolids | | Total |
|---|---|---|---|---|---|
| Sex | M | F | M | F | |
| **Grandsire family groups** | | | | | |
| AB | - | - | 4 (1) | 4 (1) | **8** |
| AC | 5 (1) (1) | 2 (1) | 1 (1) (1) | 1 (1) | **9** |
| AD | 8 (1) (1) (1) | 2 (1) (1) | 4 (1) (1) (1) | 4 (1) (1) | **18** |
| BA | 3 (1) (1) (1) | 3 (1) (1) | 2 (1) (1) (1) | 4 (1) (1) | **12** |
| BC | 4 (1) (1) | 3 (1) (1) | 2 (1) (1) | 2 (1) (1) | **11** |
| BD | 4 (1) (1) (1) | 3 (1) (1) | 6 (1) (1) (1) | 2 (1) (1) | **15** |
| CA | 2 (1) | 4 (1) | 3 (1) | 6 (1) | **15** |
| CB | 3 (1) (1) (1) | 3 (1) (1) | 3 (1) (1) (1) | 2 (1) (1) | **11** |
| CD | 6 (1) (1) (1) | 5 (1) (1) | 6 (1) (1) (1) | 6 (1) (1) | **23** |
| DA | 3 (1) (1) (1) | 1 (1) (1) | 4 (1) (1) (1) | 3 (1) (1) | **11** |
| DB | 1 (1) | 3 (1) (1) | 2 (1) (1) | 2 (1) (1) | **8** |
| DC | 1 (1) | 2 (1) | - | - | **3** |
| **Total** | **40** | **31** | **37** | **36** | **144** |

Numbers in blue parentheses depict 31 liver samples from 8-week-old F2 individuals for RRBS.
Numbers in green text depict 44 blood samples from 8-week-old F2 individuals for RRBS.
Numbers in red parentheses depict 14 sperm and seminal plasma samples from F2 for epigenetic analyses.

**Table S6.** Live born F3 progeny split by treatment, sex and great grandsire family.

| **Treatment** | **Control** | | **Biosolids** | | **Total** |
|---|---|---|---|---|---|
| **Sex** | **M** | **F** | **M** | **F** | |
| **Great grandsire family groups** | | | | | |
| ABCD | - | - | - | - | - |
| ABDC | - | - | - | - | - |
| ACBD | 3 | 2 (1) | 1 (1) | 1 | 7 |
| ACDB | 1 (1) | 1 | 3 (1) | - | 5 |
| ADBC | 3 (1) | - | 1 (1) | 2 (1) | 6 |
| ADCB | 3 (2) | 1 (1) | 4 (2) | - | 8 |
| BACD | 2 (1) | 2 (2) | 4 (1) | 5 (2) | 13 |
| BADC | 1 | - | - | - | 1 |
| BCAD | - | - | 6 | - | 6 |
| BCDA | 2 | - | 1 | 2 | 5 |
| BDAC | 2 | 2 (2) | - | 2 (2) | 6 |
| BDCA | 4 (2) | 2 (2) | 3 (2) | 1 (1) | 10 |
| CABD | - | - | - | - | - |
| CADB | - | - | - | - | - |
| CBAD | - | - | - | - | - |
| CBDA | - | - | - | - | - |
| CDAB | - | - | 3 | 2 | 5 |
| CDBA | 1(1)<br>1 | - | 2 | 1 (1) | 4 |
| DABC | - | - | - | - | - |
| DACB | - | - | - | - | - |
| DBAC | - | 1(1) | - | - | 1 |
| DBCA | - | - | - | - | - |
| DCAB | - | - | - | - | - |
| DCBA | - | - | - | - | - |
| **Total** | **22** | **11** | **28** | **16** | **77** |

Numbers in blue parentheses depict 32 available liver samples from 8-week-old F3 offspring used for RRBS.

**Dataset File S1.** Gene lists per sex and cell type for each generation.

### SI References


1. M. Bellingham et al., Exposure to a complex cocktail of environmental endocrine-disrupting compounds disturbs the kisspeptin/GPR54 system in ovine hypothalamus and pituitary gland. Environ Health Perspect 117, 1556-1562 (2009).
2. K. D. Sinclair et al., Fetal growth and development following temporary exposure of day 3 ovine embryos to an advanced uterine environment. Reprod Fertil Dev 10, 263-269 (1998).
3. N. P. Evans et al., Sexually dimorphic impact of preconceptional and gestational exposure to a real-life environmental chemical mixture (biosolids) on offspring growth dynamics and puberty in sheep. Environ Toxicol Pharmacol 102, 104257 (2023).
4. P. Dini, L. Troch, I. Lemahieu, P. Deblende, P. Daels, Validation of a portable device (iSperm(®) ) for the assessment of stallion sperm motility and concentration. Reprod Domest Anim 54, 1113-1120 (2019).
5. E. Bulkeley et al., Assessment of an iPad-based sperm motility analyzer for determination of canine sperm motility. Transl Anim Sci 5, txab066 (2021).
6. K. Gillespie, J. Stewart, Validation of iSperm analyzer for assessing ram semen quality. Proc Am Assoc Bovine Pract 58, (2024).
7. S. Parthipan et al., Spermatozoa input concentrations and RNA isolation methods on RNA yield and quality in bull (Bos taurus). Anal Biochem 482, 32-39 (2015).
8. G. W. Montgomery, J. A. Sise, Extraction of DNA from sheep white blood cells. New Zealand Journal of Agricultural Research 33, 437--441 (1990).
9. P. Di Tommaso et al., Nextflow enables reproducible computational workflows. Nat Biotechnol 35, 316-319 (2017).
10. P. Di Tommaso et al., Nextflow enables reproducible computational workflows. Nat Biotechnol 35, 316-319 (2017).
11. P. A. Ewels et al., The nf-core framework for community-curated bioinformatics pipelines. Nat Biotechnol 38, 276-278 (2020)
12. A. Kozomara, M. Birgaoanu, S. Griffiths-Jones, miRBase: from microRNA sequences to function. Nucleic Acids Res 47, D155-d162 (2019).
13. B. Langmead, C. Trapnell, M. Pop, S. L. Salzberg, Ultrafast and memory-efficient alignment of short DNA sequences to the human genome. Genome Biol 10, R25 (2009).
14. Y. Chen, L. Chen, A. T. L. Lun, P. L. Baldoni, G. K. Smyth, edgeR v4: powerful differential analysis of sequencing data with expanded functionality and improved support for small counts and larger datasets. Nucleic Acids Res 53 (2025).
15. Y. Benjamini, Y. Hochberg, Controlling the False Discovery Rate: A Practical and Powerful Approach to Multiple Testing. Journal of the Royal Statistical Society: Series B (Methodological) 57, 289-300 (1995).
16. P. Ewels, S. Peri, P. Hüther, E. Miller, N. Spix, et al., nf-core/methylseq: Endless Tofu. Zenodo. https://doi.org/10.5281/zenodo.14502249. Deposited 19 December 2024.
17. A. Mayakonda et al., Methrix: an R/Bioconductor package for systematic aggregation and analysis of bisulfite sequencing data. Bioinformatics 36, 5524-5525 (2020).
18. H. Feng, H. Wu, Differential methylation analysis for bisulfite sequencing using DSS. Quant Biol 7, 327-334 (2019).
19. G. Yu, L. G. Wang, Q. Y. He, ChIPseeker: an R/Bioconductor package for ChIP peak annotation, comparison and visualization. Bioinformatics 31, 2382-2383 (2015).
20. Q. Wang et al., Exploring Epigenomic Datasets by ChIPseeker. Curr Protoc 2, e585 (2022).
21. A. Alexa, J. Rahnenfuhrer, topGO: Enrichment Analysis for Gene Ontology. (2023) https://doi.org/10.18129/B9.bioc.topGO, R package version 2.54.0, https://bioconductor.org/packages/topGO.

22. G. Yu, L. G. Wang, Y. Han, Q. Y. He, clusterProfiler: an R package for comparing biological themes among gene clusters. Omics 16, 284-287 (2012).
23. T. Wu et al., clusterProfiler 4.0: A universal enrichment tool for interpreting omics data. Innovation (Camb) 2, 100141 (2021).
24. Q. Zhu et al., Specification and epigenomic resetting of the pig germline exhibit conservation with the human lineage. Cell Rep 34, 108735 (2021).
